# DIVE: a reference-free statistical approach to diversity-generating & mobile genetic element discovery

**DOI:** 10.1101/2022.06.13.495703

**Authors:** J. Abante, P.L. Wang, J. Salzman

**Affiliations:** Biomedical Data Science, Stanford University, 1265 Welch Rd, Palo Alto, CA 94305, USA; Center for Computational, Evolutionary and Human Genomics, Stanford University, 327 Campus Drive, Stanford, CA 94305, USA; Department of Biochemistry, Stanford University, 279 Campus Drive, Stanford, CA 94305, USA; Department of Statistics, Stanford University, 390 Serra Mall, Stanford, CA 94305, USA

**Keywords:** horizontal gene transfer, mobile genetic elements, diversity-generating mechanisms, CRISPR, transposable elements, integrative and conjugative elements, non-coding RNA

## Abstract

Diversity-generating and mobile genetic elements are paramount to microbial and viral evolution and result in evolutionary leaps conferring novel phenotypes, such as antimicrobial resistance. State-of-the-art algorithms to detect these elements have many limitations, including reliance on reference genomes, assemblers, and heuristics, resulting in computational bottlenecks and limiting the scope of biological discoveries. Here we introduce DIVE, a new reference-free approach to overcome these limitations using information contained in sequencing reads alone. We show that DIVE has improved detection power compared to existing reference-based methods using simulations and real data. We use DIVE to rediscover and characterize the activity of known and novel elements and generate new biological hypotheses about the mobilome. Using DIVE we rediscover CRISPR and identify novel repeats, and we discover unannotated genetic hyper-variability hotspots in *Escherichia coli* and *Vibrio cholerae*. Building on DIVE, we develop a reference-free framework capable of *de novo* discovery of mobile genetic elements, not currently available to our knowledge, and we use it to rediscover the known transposons in *Mycobacterium tuberculosis*, the causative agent of *tuberculosis*.

## Background

Recent estimates predict the existence of 1 trillion microbial species subject to large intraspecies variation (1, 2). This genetic diversity results from multiple processes, such as point mutations, but is increasingly appreciated to be driven by mobile genetic elements (MGEs). These elements abruptly modify the genome and rapidly provide new phenotypes (3). The rapid spread of these elements is causing a surge of antibiotic resistance, considered an urgent public health crisis (4). In addition, diversity-generating mechanisms (DGMs), such as diversity-generating retroelements and CRISPR-Cas systems, further contribute to microbial diversity. These mechanisms allow pathogens to adapt rapidly to changing environmental conditions and acquire resistance against bacteriophages. Nevertheless, current estimates point to *>* 40% of sequenced bacteria lacking annotated CRISPR systems (5), suggesting other DGMs remain to be discovered.

State-of-the-art algorithms (6–11) to detect MGEs or DGMs have many limitations, including reliance on reference genomes, assemblers, and heuristics, resulting in bottlenecks and limiting the breadth of biological discoveries. Reference genomes are only available for relatively few organisms which are, in turn, constantly evolving. Assembly algorithms are computationally expensive, especially in the metagenomic setting, and fail to capture the inherent complexity in the bacterial and viral world (12). Lastly, commonly used heuristics embedded in these algorithms impose assumptions that may not hold in general, such as the existence of target site duplications (11).

Alignment-free methods have shown great promise in several genomics problems, including transcript abundance estimation (13) and association mapping (14). However, their applications have been limited and have not been used to discover genomic variation. We reasoned that reference-free methods could have much broader discovery power in this setting. This realization led us to develop DIVE, a novel reference-free algorithm designed to identify sequences that cause genetic diversification such as transposable elements, within MGE variability hotspots, or CRISPR repeats (Fig. 1a). DIVE operates directly on sequencing reads and does not rely on a reference genome. We designed DIVE from a simple observation: a mobile genetic element such as a transposon is defined by its bounding sequence or transposon arms *A*. These arms mechanistically enable mobility, but they also define the element in an algorithmic sense: the constant sequence *A* will be flanked by a highly diverse set of sequences if *A* is indeed the arm of an MGE (Fig. 1a). This principle applies to MGE magnets and other constant/variable sequence assemblies, such as CRISPR arrays. Thus, we reasoned it should be possible to discover such elements by identifying sequences *A* with highly diverse neighboring sequences.

**Fig. 1.**
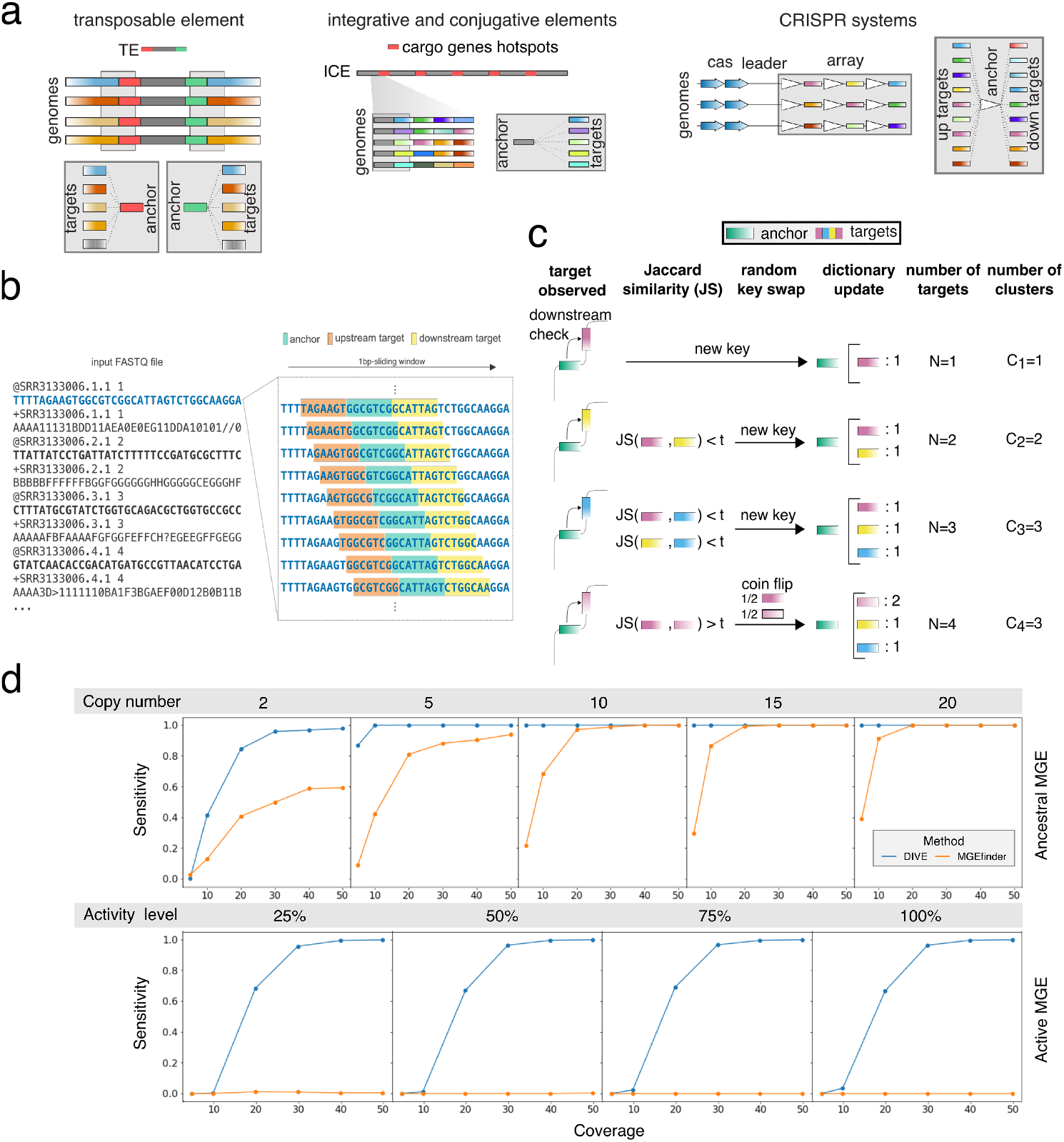
DIVE algorithm and simulations. **a**. The termini of transposable elements (TEs) shown in red (5’ end) and green (3’ end) will contain a diverse set of neighboring sequences as a result of different insertions in a genome. In turn, a highly variable set of target sequences will be observed for the anchors overlapping with the termini (red and green k-mers). Similarly, the sequences sorrounding cargo gene hotspots in integrative and conjugative elements (ICE) will be followed or preceded by a set of diverse sequences when the cargo genes vary across the analyzed sample. Lastly, CRISPR repeats will also contain a diverse set of neighboring sequences due to the diversity of spacer sequences. In this case, the diversity will be observed both upstream and downstream of the anchor as shown in the cartoon. **b**. DIVE processes reads sequentially using a sliding window that moves along the sequencing reads, recording for each anchor the upstream and downstream k-mers (targets). For each anchor, a target dictionary is constructed where DIVE keeps track of the target sequences observed clustering them as they are observed. **c**. Example of the cluster formation process based on Jaccard similarity (JS) for a given anchor (downstream case). The first target observed for the green anchor is a pink sequence. An entry is created for the green anchor in the anchor dictionary and a target dictionary is initialized for this anchor containing just the pink sequence. Then a yellow and a blue sequence are observed which, given the dissimilarity with respect to the previously observed sequences, result in a new entries in the target dictionary of the anchor. Finally, another pink sequence is observed, and given its similarity to the firstly observed pink sequence, the two are clustered together. With probability 50%, the newly observed pink sequence becomes the key in the target dictionary and the count for the pink cluster is increased. **d**. Sensitivity curves for DIVE and MGEfinder in our simulations of ancestral and active MGEs (see Supplementary Information). In the ancestral element simulations we evaluate the performance considering different copy numbers, whereas in the active element we consider various levels of element activity. The sensitivity of DIVE in detecting MGE termini is higher than that of MGEfinder in both cases for most coverages, copy number, and activity level.

DIVE makes the preceding logic into a statistical algorithm. We define k-mers *anchor* and *target*, both sequences of a predefined length *k*. DIVE aims to find anchors with neighboring (upstream or downstream) statistically highly diverse (target) sequences. DIVE processes each read sequentially using a sliding window to construct *target* dictionaries for each *anchor* encountered in each read (Fig. 1b-c). For each anchor, DIVE uses the number of target clusters *C*_*N*_, constructed using the Jaccard similarity, to compute the probability that the observed target diversity is due to background variability, resulting in a set of *p*-values that are corrected for multiple hypothesis testing (Methods). Lastly, anchors are clustered to reduce redundancies in the output, and a representative anchor (RA) maximizing the effect size *α ≡* log(*C*_*N*_ */E*[*C*_*N*_ |*N*]) in each direction is picked (Supplementary Information). Notably, *α* is a relative measure of the observed diversity compared to the expected diversity for a given coverage. This quantity can be used to prioritize CRISPR arrays with high diversity over those with less or highly mobilized over less mobile MGEs (Methods). Annotation and references are not used during any step of DIVE except as a post-facto option for interpretation. Lastly, DIVE anchors can be extended using any assembly algorithm producing longer sequences which can be subsequently used to perform protein domain homology search and classification (Methods). Together, this enables reference-free *de novo* MGE discovery, which is critical and currently unavailable to our knowledge.

## Results

### K-mer variability cannot be explained by k-mer abundance

We first evaluated the dependence of DIVE’s effect size *α* on the abundance of a k-mer in the reads by correlating it with the number of observations *N*, a proxy for the abundance of a k-mer. The lack of correlation indicates that k-mer prevalence alone does not predict diversifying sequences as expected (Fig. S1). We then used whole-genome sequence (WGS) data from *Escherichia coli* and *Vibrio cholerae* to compare the proportion of annotated RAs between DIVE and a naive algorithm that randomly picks anchors based on prevalence (Methods, Table S1, S2). RAs reported by DIVE were strongly enriched relative to the naive algorithm for mapping to known elements (Table S3), showing DIVE’s sensitivity is not a function of k-mer abundance.

### DIVE has higher sensitivity than state-of-the-art MGE termini detection methods

Next, we performed a set of simulations to systematically evaluate DIVE’s ability to identify MGE termini in two different scenarios. We considered the case where an MGE is not active and has multiple copies, and the case where a single copy of an active element has been active within a single sample (Supplementary Information; Fig. S2). DIVE achieved an area under the curve (AUC) of *>*99% in all scenarios with coverage *>* 30X (Fig. 1d, S3, S4), even when the genomic divergence in the genome was considerable. We also compared the discovery power of DIVE to that of MGEfinder, a *de novo* MGE detection tool that outperformed the state of the art in a recent benchmark (11). MGEfinder heavily relies on heuristics, an assembly step involving all the reads, and requires a reference genome, which affects the algorithm’s sensitivity (Supplementary Information). MGEfinder showed higher sensitivity than DIVE when *C*_*N*_ *≤* 5 and the mutation and indel rate (MIR) was the lowest considered. Nevertheless, DIVE showed higher sensitivity than MGEfinder in all other scenarios involving different coverages, element activity, copy number, and genome variability levels (Supplementary Information; Fig. S5, S6). We also compared the two algorithms using real *Neisseria gonorrhoeae* isolate WGS data using reads not mapping to the reference used by the authors (Supplementary Information). DIVE reported RAs mapping to over 100 known TEs (15) using data from 200 isolates (Table S4), whereas MGEfinder reported 28 unique TEs (891 isolates), consistent with the lower sensitivity observed in our simulations (Fig. 2a). In addition to known TEs, DIVE reported RAs mapping to known CRISPR direct repeats and other MGEs.

**Fig. 2.**
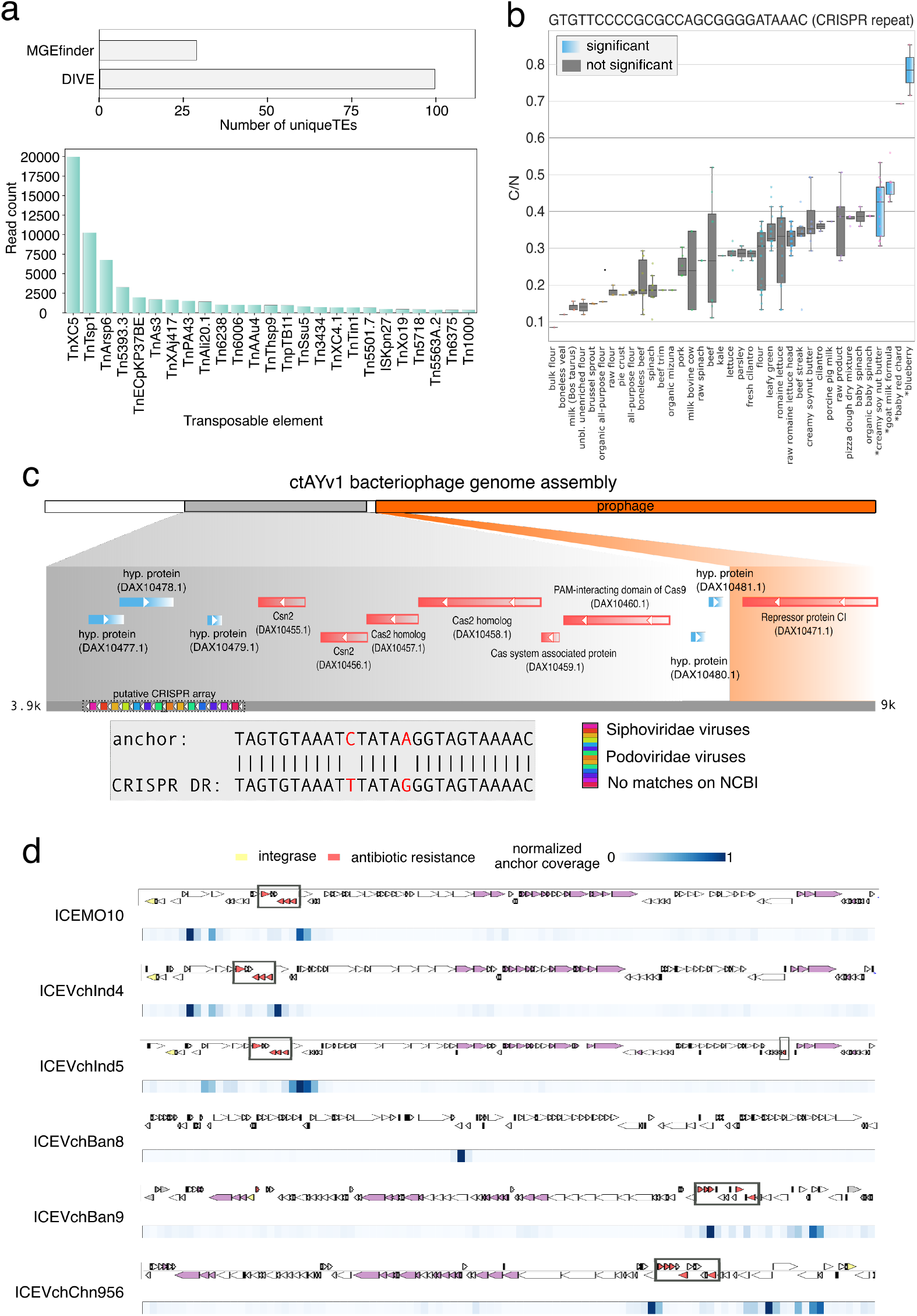
DIVE quantifies activity of mobile genetic elements and diversity-generating mechanisms. **a**. Number of distinct trans-posable elements (TEs) detected by each method and number of reads containing an anchor mapping to TE detected by DIVE among unaligned reads in *N. gonorrhoeae* data from Durrant et. al 2020. Tn3 transposons TnXc5, TnTsp1, and TnArsp6 were most prevalent, whereas TnXAj417 and Tn3434 showed the largest median effect size. **b**. Boxplot of the ratio of number of clusters to the number of observation (C/N) of the anchor sequence GTGTTCCCCGCGCCAGCGGGGATAAAC called by DIVE in *E. coli*. The target diversity in four environments (shown in blue) is significantly larger than that of the rest of environments (Methods). **c**. Putative direct repeat TAGTGTAAATCTATAAGGTAGTAAAAC detected by DIVE in the human gut metagenomic samples. The anchor reported by DIVE is two substitutions away from a known CRISPR direct repeat, and maps to an assembly (ctAYv1) containing a phage (orange) and a genomic segment (gray) having a canonical CRISPR array, derived from *Ruminococcus bromii*. **d**. Alignment of all significant anchors from the *V. cholerae* analysis to the six available sequences of integrative and conjugative elements (ICEs) in the ICEberg database. The annotated genes for each SXT ICE are shown above the plot (yellow genes: integrases; red genes: antibiotic resistance). Below the ICE, a heatmap shows the coverage of anchors with respect to the element. Anchors cluster in the neighborhood of known antibiotic resistance genes in all but ICEVchBan8, where they overlap with a known transposase.

### De novo detection of CRISPR repeats

We next tested whether DIVE could identify CRISPR repeats *de novo* in *E. coli* using purely statistical features. Three hundred one of the RAs identified by DIVE mapped to known CRISPR repeats, with their associated targets mapping to 774 known spacers in the CRISPR-Cas++ database (8) (Methods). In particular, of the 59 known *E. coli* repeats present in the data and meeting the minimum sample size (*N*_min_ = 25), DIVE detected 50 (85%). Additionally, DIVE detected eight repeats not annotated in *E. coli* (seven known in Enterobacteria and one in Proteobacteria) (Table S5). Of all categories, RAs mapping to CRISPR repeats showed the largest median effect size (Fig. S1). They were further distinguishable from other RAs in that they belonged to anchor clusters with bidirectional significance, showed large effect size (0.5(*α*_*u*_ + *α*_*d*_) *≥* 3.5) and appeared more than once within a read. Imposing these requirements restricted to ten RAs, perfectly mapping to six known *E. coli* CRISPR repeats (Table S6), achieving 100% specificity. This observation suggests that these criteria may be used to identify CRISPR repeats *de novo*. Using these criteria, we identified a novel CRISPR repeat in human gut metagenomic samples (Fig. 2c). This repeat was related but not identical to a known repeat in the CRISPR-Cas++ database, and formed an array with many spacers mapping to known phages. BLAST of the RA to the NCBI WGS database restricted to the bacterium *Ruminococcus bromii* gave perfect matches in a locus with CRISPR-associated (Cas) genes, suggesting this RA constitutes a novel functional CRISPR DR (Supplementary Information). We also used these criteria to identify six novel repeats in microbiota metagenomic samples (Methods). This set of repeats included five with 1-5 mismatches to the closest repeat on CRISPR-Cas++ and one that did not match any known repeat (Fig. S7). Three of the six repeats completely aligned to at least one bacterial genome on NCBI (BLAST). These repeats had varying copy numbers in the genomes, ranging from 1 to 12. In addition, we found annotated CRISPR *Cas* and *Csd* proteins nearby for two of the repeats. Interestingly, the array consisted of just 1-2 repeats in these cases, constituting mini-CRISPR arrays. This result shows that DIVE permits the discovery of CRISPR arrays lacking *Cas* genes in *cis* since it does not rely on heuristics such as nearby known *Cas* proteins, as well as arrays with low numbers of repeats. DIVE provides additional information about CRISPR repeats beyond discovering them *de novo*, establishing a framework to estimate spacer diversity in CRISPR arrays as a function of available covariates when multiple samples are available. The target clusters produced by DIVE for a given anchor can be compared across different environments or conditions and statistically modeled using a binomial generalized linear model to identify covariates driving the differences in target diversity (Methods). In the *E. coli* dataset, five RAs mapped to *E. coli* CRISPR repeats with spacer (target) diversity being a function of the isolation source (Fig. 2b, S8, S9). In addition to CRISPR, several other loci had high target diversity, including 77 RAs mapping to lambdoid phages, ICEs, TEs, and non-coding RNA (ncRNA), where target diversity varied as a function of the isolation source (Fig. S8, S10, S11). Thus, this framework can enable the study of genetic diversity as a function of available covariates beyond CRISPR arrays.

### Rediscovery of non-coding RNA as MGE insertion hotspots

Next, we investigated whether DIVE could rediscover the role that non-coding RNA (ncRNA) loci play in the genome as insertion hotspots for MGEs (16, 17) (Supplementary Information). For example, regions coding for tRNA genes are common insertion sites for phages and plasmids (18). DIVE found substantial target diversity among RAs mapping to ncRNAs, including several examples where the target variability was a function of available covariates (Fig. S11). In *E. coli*, DIVE called 712 unique RAs mapping to 84 different Rfam accessions annotated as tRNA genes (BLAST *e*<0.01), aligning to the ends of the genes (Table S7). DIVE RAs also mapped to 82 Rfam accessions annotated as nucleoid-associated noncoding RNA 4 (naRNA4) genes (Table S8). These genes are encoded in repetitive extragenic palindrome (REP) regions, which have been previously observed at the recombination junctions of lambda phages and described as hotspots for transposition events (19, 20). DIVE also identified 264 RAs mapping (BLAST *e*<0.01) to 26 Rfam accessions annotated as antisense RNA 5 (asRNA5) among *V. cholerae* isolates, four of which have a target diversity significantly associated with available covariates, including date, country, and isolation environment (Table S9).

### A reference-free framework for MGE *de novo* discovery

DIVE can detect many kinds of sequence diversity. To specifically illustrate DIVE’s application to reference-free *de novo* MGE discovery, we analyzed a subsample of seventy-five *M. tuberculosis* isolates from the CRyPTIC consortium (21) and focused on *de novo* discovery of transposon sequences. Using the RAs reported by DIVE for each isolate, we reconstructed putative transposons by performing seed-based assembly on the RAs showing hyper-variability in a single direction since that is the expected behavior in MGE termini. Subsequently, we performed protein homology analysis on the resulting contigs, after translating them *in silico*, allowing us to identify sequences with transposon-associated domains for each isolate (Fig. 3a; Methods). We clustered the resulting contigs using CD-HIT-EST (22) to collapse similar sequences from different isolates (>90% identity), producing 47 clusters (of which 37 were singletons) with sizes ranging from 236 to 20,000 bp. The contig assemblies showed good fidelity, as all BLAST alignments of the longest contig from each cluster gave near full-length matches in genomes from *M. Tuberculosis* complex with *≥*99% identity. Furthermore, we observed different degrees of alignment multiplicity across the contigs, which we attribute to differences in copy number (Fig. 3b; Methods). Alignment to the database ISfinder (23) showed that 19 clusters contained strong matches to insertion sequences (BLAST *e ≤*2e-75; Table S10). Nine clusters matched IS*6110* (BLAST *e* =0), a well-known insertion sequence in *M. Tuberculosis* commonly used to type strains (Fig. 3d). When we aligned the input seeds in the assembly step onto this element, we found they accumulated on the ends as expected (Fig. 3c). Eleven other clusters matched nine known transposon sequences in *Mycobacteria*, among which we completely assembled the sequences of IS*Mt1* (Fig. 3e), IS*Mt2*, IS*Mt3*, IS*1553* and IS*1538* (Table S10). Among the remaining 28 clusters, BLAST of representative contigs against the NCBI database showed that 14 could be explained by other repetitive elements (lying near to insertion sequences), the major categories being PE/PPE genes (24) and REP13E12/HNH endonuclease/DUF222 (25, 26) (Table S10). To our knowledge, such a reference-free approach for *de novo* discovery of MGEs is not currently available.

**Fig. 3.**
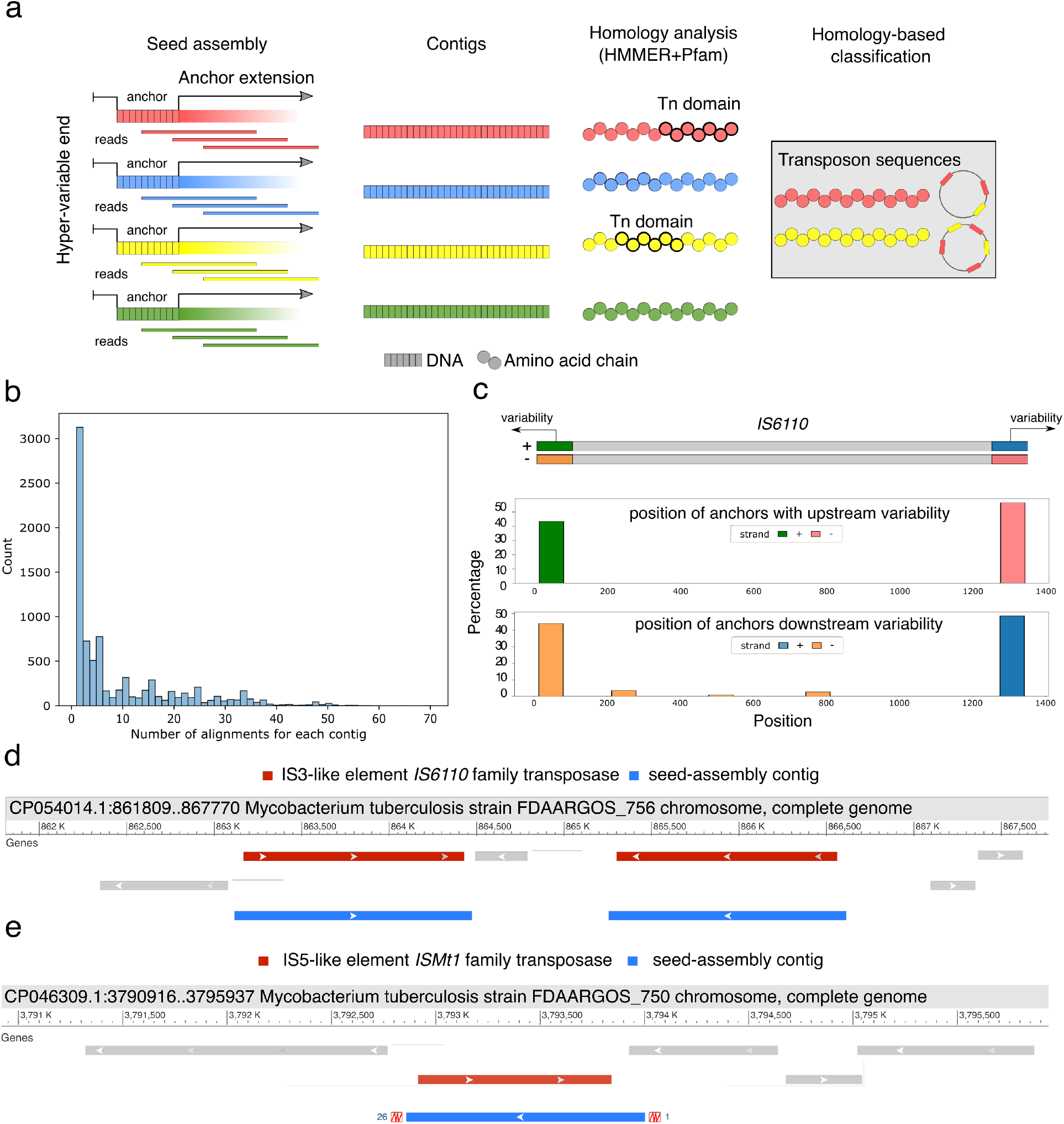
De novo discovery of transposons. **a**. DIVE anchors with variability in a single direction are chosen as putative transposon termini that can be used as seeds in an seeded assembly step. The resulting contigs are translated *in silico* to produce amino acid sequences. Lastly, protein domain homology can be performed on the resulting protein sequences to identify putative transposons, as well as other elements of interest. **b**. Histogram of the number of alignment loci for each contig in at least a *M. tuberculosis* genome in the NCBI nucleotide database. **c**. Pile up of anchors in IS*6110* concentrating on the ends of the insertion sequence, which has a length of 1354 bp. **d**. BLAST alignment of contig104 against a *M. tuberculosis* genome, mapping to two copies of the element inserted in opposite directions. This contig corresponds to the insertion sequence IS*6110*. **e**. BLAST alignment of contig2 against a *M. tuberculosis* genome. This contig corresponds to the insertion sequence IS*Mt1*.

### Evidence of extensive sequence diversity in MGE cargo

State-of-the-art methods do not attempt to discover within-MGE variability, which is known to have a significant phenotypic impact (27). We hypothesized that DIVE could discover hotspots of genetic rearrangements within MGEs. As an example, integrative and conjugative elements (ICEs) confer various properties to their hosts through cargo genes, such as phage and antibiotic resistance (28). In *V. cholerae*, SXT ICEs determine phage resistance, and deletion of hotspot five in SXT ICEs leads to susceptibility to ICP1 phage infection (27). We hypothesized that DIVE could recognize the boundaries of cargo gene hotspots in SXT ICEs in *V. cholerae* isolates. To test this hypothesis, we looked at the concentration of anchors around known SXT cargo hotspots. DIVE’s RAs clustered around the edges of cargo gene hotspots in five SXT ICE variants and a putative hotspot in an unannotated SXT ICE (Fig. 2d, Supplementary Information). Extension of these anchors through an assembly algorithm could enable the study of differences in antibiotic resistance gene composition.

### Unannotated genetic hyper-variability in *E. coli* and *V. cholerae*

DIVE also discovered a large number of unannotated genetic hyper-variable regions in well-studied organisms like *E. coli* and *V. cholerae*. We filtered the unannotated RAs with at least a median *α ≥* 2 and a sample prevalence *>* 5%, resulting in a shortlist of 5 anchors among *E. coli* isolates and 29 among *V. cholerae* isolates (Table S11), and here we highlight two examples (Supplementary Information). In each case, raw reads containing the corresponding RA were multiple-sequence aligned (29) showing a sudden decay in consensus nucleotide composition on one end of the RA consistent with DIVE’s output. In the *E. Coli* example, the RA produces a unique match in the *E. coli* O157 reference proximal to the 3’ end of isomerase Arabinose 5-phosphate isomerase (kdsD), an intramolecular oxidoreductase that interconverts aldoses and ketoses, with target diversity that results in downstream non-synonymous mutations and indels in the enzyme (Fig. 4a). In *V. cholerae*, the RA produced a single match in at least one hundred NCBI *V. cholerae* accessions, mapping to the intergenic region between tRNA-Cys and the artP gene (arginine ABC transporter ATPase), with the former being in the direction of more target diversity. The target diversity observed was not well represented on the reference genomes available on NCBI, with *>* 30% of targets not blasting to any genome in either case (Fig. 4b). Furthermore, the sequence variability extended well beyond the range of the target sequence, spanning over 100 nt, including substitutions and indels altering the protein sequence. These unannotated hyper-variable regions warrant further investigation to identify the mechanism and functional consequences of their sequence variability.

**Fig. 4.**
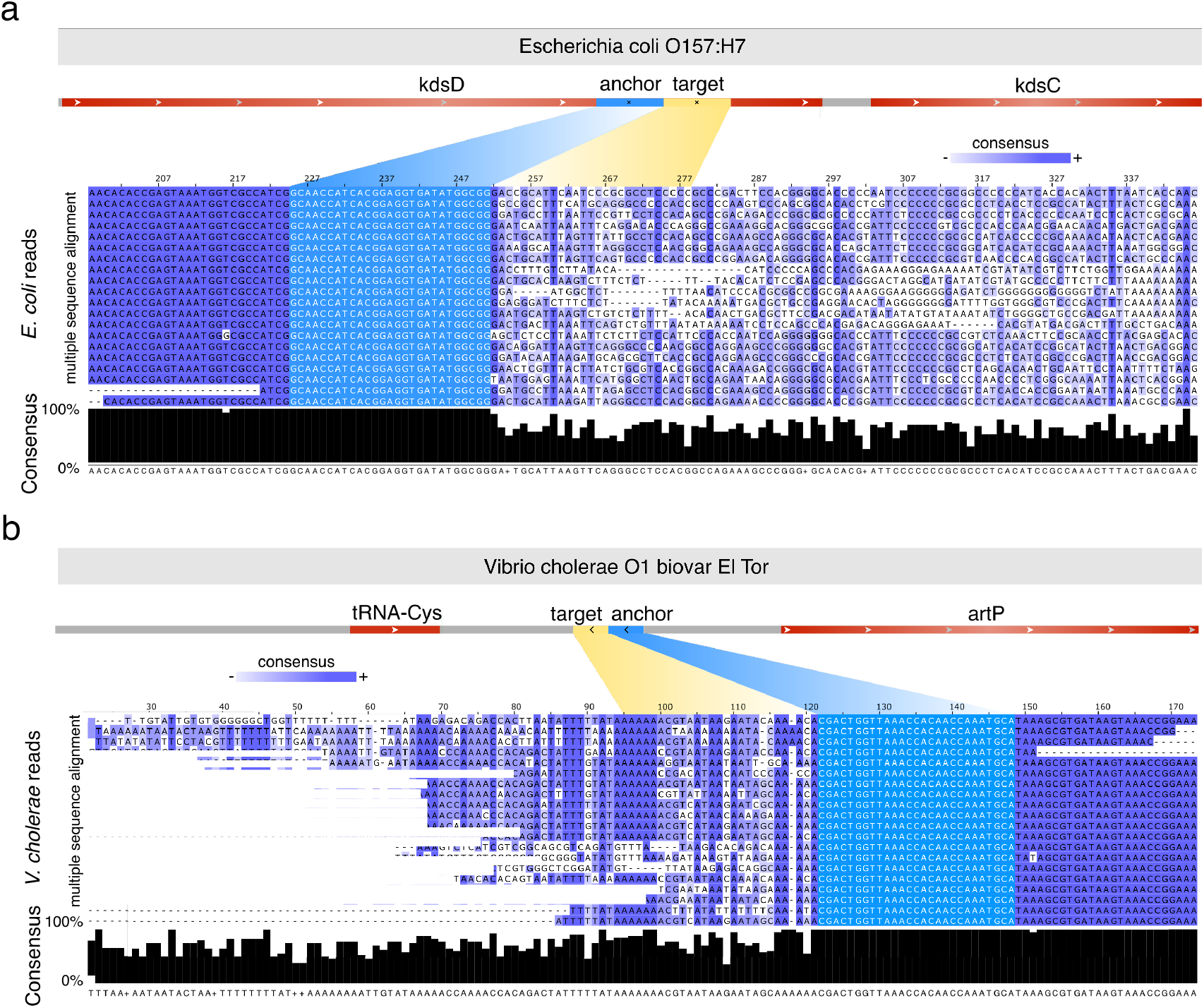
DIVE discovers unexplained hyper-variability in *E. coli* and *V. cholerae*. **a**. The anchor CCGCCATATCACCTCCGTGATG-GTTGC showed the largest median effect among unannotated anchors in *E. coli* (*α*=4.22, prevalence=5.28%). The anchor produces a unique match in *E. coli* O157 overlapping isomerase Arabinose 5-phosphate isomerase (kdsD). The multiple sequence alignment (CLUSTALW) of a random selection of 20 reads containing the unannotated anchor (below). The sequence consensus (black bars) is approximately 100% upstream of the anchor (light blue), whereas downstream of the anchor the sequences found across the reads diverge significantly and introduce substitutions and indels. **b**. The unannotated anchor CCGCCATATCACCTCCGTGATGGTTGC showed the largest median effect size among unannotated anchors in *V. cholerae* (*α*=3.47, prevalence=5.17%). The anchor produced a single match in at least one hundred NCBI accessions, all *V. cholerae* strains, mapping to the intergenic region between tRNA-Cys and the artP gene (arginine ABC transporter ATPase), with the former being in the direction of more target diversity. The multiple sequence alignment (CLUSTALW) of a random selection of 20 reads containing the unannotated anchor (below). The sequence consensus (black bars) is approximately 100% upstream of the anchor (light blue), whereas downstream of the anchor the sequences found across the reads diverge significantly and introduce substitutions and indels.

## Discussion

DGMs and MGEs are crucial to microbial evolution, driving evolutionary leaps and leading to the emergence of novel phenotypes such as antimicrobial resistance (3). However, detecting these elements remains a challenge, as existing algorithms have several limitations such as reliance on reference genomes, assemblers, and heuristics, resulting in computational bottlenecks and limiting the scope of biological discoveries (12).

To overcome these limitations, we propose a new reference-free approach that uses information contained in sequencing reads alone and we demonstrate that this approach has improved detection power compared to state-of-the-art reference-based methods, such as MGEfinder (11), both through simulations and real data. We show that DIVE can detect MGE termini both for ancient and active elements. Moreover, DIVE can perform well even when there is important sequence divergence within the sample analyzed, in contrast to the state of the art. This is mainly due to the fact that DIVE can detect elements without relying on a reference genome, making it a valuable especially for organisms with no reference genome.

DIVE can be used to discover DGMs *de novo*. Here we used DIVE to rediscover CRISPR and identify novel repeats. In particular, in *E. coli*, we were able to rediscover 85% of the repeats present in the data, and we discovered several novel repeats in metagenomic samples, including two constituting mini-CRISPR arrays. Furthermore, we illustrate how DIVE’s output can be used to establish associations between sequence variability and available covariate information. For example, we identified five CRISPR repeats for which their spacer variability is a function of the isolation source in *E. coli*, and we identified other elements for which the variability was associated to the isolation date and country in *V. cholerae*. Using DIVE, we also rediscovered ncRNA coding regions as MGE insertion hotspots (16, 17), including tRNA and naRNA4, in*E. coli* and *V. cholerae* isolates. We also found asRNA5 loci to show hyper-variability in *V. cholerae*, in some instances associated to the isolation date, country, and environment.

Combining DIVE with a seed-based assembler and a protein domain homology algorithm, we propose a reference-free framework for *de novo* discovery of MGEs, which to the best of out knowledge is not currently available. We used this framework to rediscover the majority of known insertion sequences in *M. tuberculosis*. This framework could also be used to discover other types of MGEs, for example phages, by filtering the contigs for phage-associated protein domains. We also show how DIVE can be used to study MGE intra-variability in the context of SXT ICE elements in *V. cholerae*, which determinines phage resistance (27), where we rediscover cargo gene hotspots boundaries. Extending the target sequences in this case would enable further analysis of the relationship between cargo gene content and antibiotic resistance phenotypes.

Lastly, DIVE can discover unannotated genetic hyper-variability hotspots in genomes. Here we identify several loci in *E. coli* and *V. cholerae* presenting a sudden decay in sequence consensus. Furthermore, we highlight two examples where the decay occurs in coding regions leading to protein sequence alterations. Thus, we believe these loci warrant further investigation to identify the mechanism and functional consequences of their sequence variability.

A current limitation of DIVE is that it is designed to analyze a single sample at a time. Nevertheless, in certain contexts, it might be of interest to study differential variability across groups, for example (30). Furthermore, we envision that the assembly process could be better focused by leveraging information from other anchors. For example, the extension process could stop when another anchor, for which we observed downstream hyper-variability, is included in the contig.

Overall, we believe the development of DIVE provides a valuable contribution to the field of microbial genomics. The ability to detect DGMs and MGEs without relying on reference genomes, and the development of a *de novo* discovery framework for MGEs, expands the scope of biological discoveries that can be made.

## Conclusions

DIVE offers a new reference-free paradigm for studying genetic bacterial evolution. It is a statistically-grounded algorithm, providing probabilities that can be used to assess the statistical significance of the discoveries and to control for false positives. DIVE offers higher sensitivity than state of the art methods both in simulations and real data. In addition, DIVE is a reference-free algorithm, which makes it an ideal candidate to study DGMs and MGEs in settings where a reference genome is not available. Furthermore, DIVE can be used in conjunction with seed-based assemblers and protein domain homology algorithms, enabling reference-free *de novo* discovery of MGEs. More importantly, given the generality of the principle, DIVE could enable the discovery of entirely novel biological mechanisms that diversify genomes. Overall, DIVE offers numerous new possibilities and research avenues to study bacterial evolution.

## Methods

### DIVE configuration

Here we defined the k-mer size *k* to be 27, striking a balance between the memory requirements and the specificity of the sequence. DIVE also allows the user to introduce a gap in between the anchor and the targets of an arbitrary distance *g*, which here we set at *g* = 0. We defined the minimum and the maximum number of targets per anchor to be *N*_min_ = 25 and *N*_max_ = 75, respectively. Finally, to strike a balance between power and computational resources, we decided to limit the number of FASTQ records processed per sample to 2.5M.

### Clustering targets on the fly

Upon observing a target k-mer, DIVE checks whether the k-mer produces a Jaccard similarity (JS) larger than a given threshold (0.2) with any of the previously observed targets. If so, the counter of that key is increased by one, and, with a probability of 50%, the key is replaced by the newly observed target sequence. If no such target sequence exists in the dictionary, a new key is created, and its counter is initialized at one. To calculate the JS between two targets of length *k* = 27, DIVE splits each one into smaller k-mers of length seven and computes the JS similarity between the two using

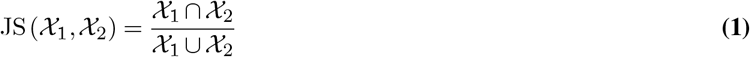

where *X*_*i*_ is the set of 7-mers in the *i*-th target sequence.

### Minimum and maximum N

For each anchor, DIVE requires a minimum number of targets observed *N*_min_ to proceed with the downstream statistical analysis. In addition, since it might not be necessary to keep all the targets observed exhaustively, DIVE also imposes a maximum number of targets *N*_max_ to be observed for each anchor. This allows the algorithm to skip anchors for which it has already observed *N*_max_ targets in each direction and move faster along the FASTQ file. Here we set these parameters to *N*_min_ = 25 and *N*_max_ = 75.

### Minimum sample size prediction

To manage the memory burden and make the algorithm more efficient, we impose a minimum number *N*_min_ of targets to be observed by the end of the FASTQ file. We let the total number of FASTQ records in the file be *L* and the number of observed records so far be *l*. Then, letting *p*_min_=*N*_min_*/L* be the probability that we will observe a target sequence for any given anchor in a single read at least once, we compute the probability that, after *l* records observed, we have observed zero instances using

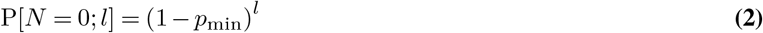

When this probability becomes smaller than 0.01 this implies that the rate we are observing targets for the anchor is too slow for us to observe *N*_min_ by the end with 99% probability. Thus, DIVE does not accept new anchors into the anchor dictionary DA past this point. This condition allows us to discard new anchors that appear late in the FASTQ file, which will likely not produce enough data to test for hyper-variability, reducing an unnecessary burden for dictionary *D*_*A*_. Past the point where DIVE does not accept new anchors, DIVE also anticipates the number of targets that will be observed for each anchor remaining in the dictionary to decide whether a given anchor should remain in the dictionary *D*_*A*_. More precisely, the oracle used by DIVE computes the probability that we will observe at least *N*_min_ targets by the end of the FASTQ file. Letting *L* be the total number of FASTQ reads, *x* the number of targets observed, and *l* the number of FASTQ reads processed so far, we can compute

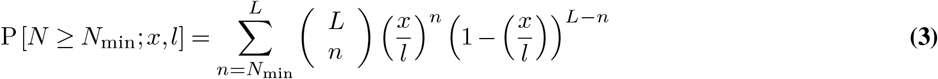

where *x/l* is an empirical estimate of the probability of observing a target sequence for the given anchor at least once in a single read. However, we use the normal approximation to the binomial distribution for efficiency to compute this probability. The anchor is conserved in the dictionary *D*_*A*_ if this probability is larger than 50%. Otherwise, the anchor is removed to reduce the size of the dictionary and speed up the algorithm.

### Target sequence hyper-variability

We let *X*_*n*_ be a binary random variable indicating whether we created a new key in the target sequence dictionary upon observing the *n*-th target sequence for a given anchor and direction. Let *C*_*n*_ be the total number of keys target sequence clusters (keys) formed after observing the *n*-th target sequence. Then, assuming that *X*_*n*_ is independent and distributed according to a Bernoulli distribution with success probability *p*_*n*_, we have that *C*_*N*_ *∼* PoissBin(*p*_1_, …, *p*_*N*_). Upon observing a set of trajectories *{C*_1_, *C*_2_, …, *C*_*M*_ *}*, the success probabilities *{p*_*n*_*}* can be estimated from the data by simply computing

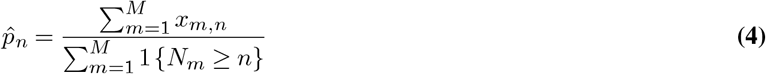

This follows from the fact that we can assume that most of the anchors will not be nearby hyper-variable regions. Nevertheless, the following test will be more conservative if we happen to include anchors adjacent to hyper-variable regions by chance. For a given *C*_*N*_ = *c*, we can compute the probability under the null hypothesis that the observed value *c* can be explained by the background biological (e.g., point mutations) and technical variability (e.g., sequencing errors) using

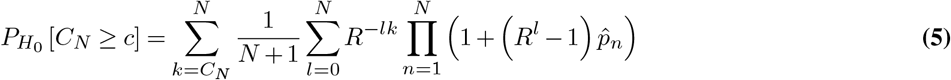

Note that the JS threshold determines the propensity of creating new target clusters. However, the null distribution of this test statistic is estimated from the dataset and thus is created for the specific JS threshold chosen by the user and the test will be valid regardless of the threshold. Nevertheless, this threshold needs to be small enough so that whenever there is true diversity new target clusters are formed so that differences can be observed. At the same time, having a very high JS threshold would result in very large target dictionaries since for every new sequence observed with a single nucleotide difference a new target cluster would be created. Even though the hypothesis testing framework could handle this situation, this would require a lot of RAM memory since the target dictionary of every single anchor would grow substantially. In our work we found that a 0.2 threshold achieves a good compromise for *k*=27.

The resulting p-values are corrected using the Benjamini-Hochberg (BH) correction, resulting in a list of q-values that allow us to perform false discovery rate (FDR) control.

### Efficient Benjamini-Hochberg correction

To correct for multiple hypothesis testing, we DIVE implements a memory-efficient version of the Benjamini-Hochberg (BH) correction. DIVE records the total number of statistical tests performed *m* and drops non-significant cases (*p >* 0.1). Then, it adjusts the remaining *m′* p-values by using the following procedure. First, it ranks the remaining *m′ p*-values *{p*_*i*_*}* in ascending order *{p*^(1)^, *p*^(2)^, …, *p*^(*m′*^)*}*. Then, it computes the adjusted p-value using *q*^(*i*)^ = *p*^(*i*)^*m/i*, and it reports the anchors for which *q <* 0.1. Note that p-values computed for overlapping k-mers will be positively correlated. Nevertheless, the BH correction can still control the FDR under that form of dependence (31).

### Clustering of anchors

Once the BH correction is applied to the computed p-values, DIVE uses DBSCAN (Levenshtein edit distance) to cluster the anchor sequences to avoid redundancies in the output. This is done using the function DBSCAN from the python package sklearn with parameters eps=2 and min_samples=1. For each anchor cluster, a representative anchor is chosen for each direction (upstream and downstream) by picking the anchor that maximizes the effect size.

### Effect size computation

The effect size is quantified using the log-fold change between the observed number of clusters *c*_*N*_ and the expected number of clusters for a given number of observed targets, as quantified by *N*, and it is given by

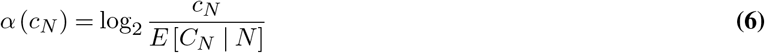

This quantity can be used to rank the results in terms of effect size and allows the user to filter out positives with small effect size and thus little biological interest.

### Annotation of DIVE’s output

To validate our results, we used blastn-short 2.11.0 (32) to align the detected k-mers to a set of sequence databases stored in FASTA format comprised of CRISPR direct repeats (CRISPR-Cas++ (8)), transposable elements (Dfam (33); TnCentral (15)), mobile genetic elements (ACLAME (34)), internal transcribed spacers 1 and 2 (ITSoneDB (35), ITS2 (36)), integrative and conjugative elements (ICEberg (37)), and RNA sequence families of structural RNAs (Rfam (38)). Furthermore, we cross-referenced our results with Illumina adapter (obtained from TrimGalore) and UniVec sequences (NCBI) to remove technical artifacts. We classified each representative anchor (upstream and downstream) as unannotated if no BLAST produced *e ≤* 0.25, questionable if 0.01 *≤ e <* 0.25, and annotated if the lowest *e ≤* 0.01 with the corresponding annotation. Here we chose this comprehensive set of annotation files. Nevertheless, DIVE can take an arbitrary number of annotation files as input, and thus this step can be adjusted depending on the application.

### Enrichment analysis of positive controls

A total of 100,000 27-mers were chosen at random among the entire *E. coli* dataset, allowing for repetition. We used BLAST to align this set of random anchors to the set of annotation files we used during our analysis. Then, we deemed anchors as unannotated and annotated if their BLAST e-values were *e>0*.*25* and *e<0*.*01*, respectively. Then, we used the function fisher_exact python’s scipy package to perform a two-sided Fisher’s exact test comparing the proportions of annotated to unannotated between this set of random anchors and the set of DIVE anchors.

### Binomial regression

To find associations between the target variability and a set of covariates given by **x**, we use a binomial regression model such that *C ∼* Binomial(*N* ; *p*(**x**)), such that

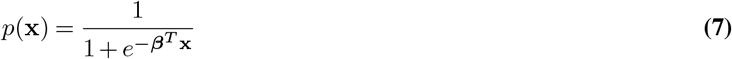

For *E. coli* isolates, we used the isolation source as a covariate, whereas for *V. cholerae* isolates we used the environment, date, country, as well as first-order interactions.

### A posteriori clustering of targets

To cluster the targets across samples *a posteriori*, we used a greedy clustering approach. We rank the targets based on their total count across the entire dataset. Starting from the most prevalent target, we recruit all targets for which the JS>0.2 and eliminate these from the list. We proceed with the most prevalent target sequence remaining in the list the same way, and we repeat this procedure until no sequences are left. The number of times we repeat this procedure defines the number of across-sample target sequence clusters.

### Anchor extension and protein domain homology

In a given sample, DIVE anchors can be extended using an assembly algorithm to produce a longer sequence of interest. To that end, we used SSAKE (v4.0) in the seed mode iteratively (-p 1 -m 20 -o 1 -c 1 -w 5 -r 0.51 -i 0). In each iteration, SSAKE produced a set of contigs that where used in the next iteration as seeds. In our analysis, we performed ten iterations. This process results in a set of extended anchors that we subsequently translate *in silico*, resulting in a set of amino acid sequences that can be used as input to protein domain homology search algorithms. To that end, we used HMMER (v3.3.2) in conjunction with Pfam35 in the default setting. In our example, we search for transposon-associated domains since we are interested specifically in the detection of such elements. However, the list of domains is application-specific and other elements could be prioritized by using the corresponding protein domain list.

## Declarations

### Availability of data and materials

The data sets analyzed in this paper are publicly available and published. The *E. coli* isolate sequencing data used was generated by the GenomeTrakr Project, US Food and Drug Administration (PRJNA230969). We used 514 samples encompassing 62 different isolation sources (Table S12). The *V. cholerae* sequencing data was downloaded from SRA (PRJNA723557). We used all the 247 *V. cholerae* isolates sequenced as part of this project (Table S13). The rotavirus metagenomics sequencing data was downloaded from SRA (PRJNA729919). We used 102 in our analysis (Table S14). Finally, the *N. gonorrhoeae* data was downloaded from SRA (PRJNA298332), from which we used 200 samples in our analysis (Table S14).

## Code availability

The python package biodive developed in this work is publicly available at https://github.com/jordiabante/biodive. The code used to analyze the output of DIVE is publicly available at https://github.com/jordiabante/DIVEpaper.

## Authors’ contributions

J.A. and J.S. developed the computational method. J.A. developed the software. J.A., P.W. and J.S. analyzed the data. J.S. conceived and supervised the project. J.A. and J.S. wrote the paper with the assistance of P.W..

## Acknowledgements

We would like to thank Ruth Timme, David Lipman, Andrew Fire, Kaitlin Chuang, Roozbeh Dehghannasiri, Elisabeth Meyer, Tavor Baharav and Ivan Zheludev for their comments and suggestions. We would also like to thank Kaitlin Chuang for her help processing part of the data used in this paper.

## Bibliography

1. Kenneth J Locey and Jay T Lennon. Scaling laws predict global microbial diversity. Proceedings of the National Academy of Sciences, 113(21): 5970–5975, 2016.

2. Ruiting Lan and Peter R Reeves. Intraspecies variation in bacterial genomes: the need for a species genome concept. Trends in microbiology, 8 (9):396–401, 2000.

3. Chitra Dutta and Archana Pan. Horizontal gene transfer and bacterial diversity. Journal of biosciences, 27(1):27–33, 2002.

4. Rachel A Smith, Nkuchia M M’ikanatha, and Andrew F Read. Antibiotic resistance: a primer and call to action. Health communication, 30(3): 309–314, 2015.

5. Rotem Sorek, Victor Kunin, and Philip Hugenholtz. Crispr—a widespread system that provides acquired resistance against phages in bacteria and archaea. Nature Reviews Microbiology, 6(3):181–186, 2008.

6. Aaron E Darling, Bob Mau, and Nicole T Perna. progressivemauve: multiple genome alignment with gene gain, loss and rearrangement. PloS one, 5(6):e11147, 2010.

7. Abraham G Moller and Chun Liang. Metacrast: reference-guided extraction of crispr spacers from unassembled metagenomes. PeerJ, 5:e3788, 2017.

8. David Couvin, Aude Bernheim, Claire Toffano-Nioche, Marie Touchon, Juraj Michalik, Bertrand Néron, Eduardo PC Rocha, Gilles Vergnaud, Daniel Gautheret, and Christine Pourcel. Crisprcasfinder, an update of crisrfinder, includes a portable version, enhanced performance and integrates search for cas proteins. Nucleic acids research, 46(W1):W246–W251, 2018.

9. Panisa Treepong, Christophe Guyeux, Alexandre Meunier, Charlotte Couchoud, Didier Hocquet, and Benoit Valot. panisa: ab initio detection of insertion sequences in bacterial genomes from short read sequence data. Bioinformatics, 34(22):3795–3800, 2018.

10. Alexander Mitrofanov, Omer S Alkhnbashi, Sergey A Shmakov, Kira S Makarova, Eugene V Koonin, and Rolf Backofen. Crispridentify: identification of crispr arrays using machine learning approach. Nucleic acids research, 49(4):e20–e20, 2021.

11. Matthew G Durrant, Michelle M Li, Benjamin A Siranosian, Stephen B Montgomery, and Ami S Bhatt. A bioinformatic analysis of integrative mobile genetic elements highlights their role in bacterial adaptation. Cell host & microbe, 27(1):140–153, 2020.

12. Izaak Coleman and Tal Korem. Embracing metagenomic complexity with a genome-free approach. Msystems, 6(4):e00816–21, 2021.

13. Rob Patro, Stephen M Mount, and Carl Kingsford. Sailfish enables alignment-free isoform quantification from rna-seq reads using lightweight algorithms. Nature biotechnology, 32(5):462–464, 2014.

14. Atif Rahman, Ingileif Hallgrímsdóttir, Michael Eisen, and Lior Pachter. Association mapping from sequencing reads using k-mers. Elife, 7:e32920, 2018.

15. Karen Ross, Alessandro M Varani, Erik Snesrud, Hongzhan Huang, Danillo Oliveira Alvarenga, Jian Zhang, Cathy Wu, Patrick McGann, and Mick Chandler. Tncentral: a prokaryotic transposable element database and web portal for transposon analysis. MBio, 12(5):e02060–21, 2021.

16. Rolf Marschalek, Thomas Brechner, Elfi Amon-Böhm, and Theodor Dingermann. Transfer rna genes: landmarks for integration of mobile genetic elements in dictyostelium discoideum. Science, 244(4911):1493–1496, 1989.

17. Thomas H Eickbush and Danna G Eickbush. Finely orchestrated movements: evolution of the ribosomal rna genes. Genetics, 175(2):477–485, 2007.

18. Allan M Campbell. Chromosomal insertion sites for phages and plasmids. Journal of bacteriology, 174(23):7495–7499, 1992.

19. Michiyo Kumagai and Hideo Ikeda. Molecular analysis of the recombination junctions of λ bio transducing phases. Molecular and General Genetics MGG, 230(1):60–64, 1991.

20. Raquel Tobes and Eduardo Pareja. Bacterial repetitive extragenic palindromic sequences are dna targets for insertion sequence elements. BMC genomics, 7(1):1–12, 2006.

21. Timothy M Walker, Paolo Miotto, Claudio U Köser, Philip W Fowler, Jeff Knaggs, Zamin Iqbal, Martin Hunt, Leonid Chindelevitch, Maha R Farhat, Daniela Maria Cirillo, et al. The 2021 who catalogue of mycobacterium tuberculosis complex mutations associated with drug resistance: a genotypic analysis. The Lancet Microbe, 3(4):e265–e273, 2022.

22. Limin Fu, Beifang Niu, Zhengwei Zhu, Sitao Wu, and Weizhong Li. Cd-hit: accelerated for clustering the next-generation sequencing data. Bioinformatics, 28(23):3150–3152, 2012.

23. Patricia Siguier, Jocelyne Pérochon, L Lestrade, Jacques Mahillon, and Michael Chandler. Isfinder: the reference centre for bacterial insertion sequences. Nucleic acids research, 34(suppl_1):D32–D36, 2006.

24. Christopher D’Souza, Uday Kishore, and Anthony G Tsolaki. The pe-ppe family of mycobacterium tuberculosis: Proteins in disguise. Immunobiology, 228(2):152321, 2023.

25. Stephen V Gordon, Beate Heym, Julian Parkhill, Bart Barrell, and Stewart T Cole. New insertion sequences and a novel repeated sequence in the genome of mycobacterium tuberculosis h37rv. Microbiology, 145(4):881–892, 1999.

26. Conserved protein domain family DUF222. https://www.ncbi.nlm.nih.gov/Structure/cdd/PF02720. Accessed: 2023-05-04.

27. Kristen N LeGault, Stephanie G Hays, Angus Angermeyer, Amelia C McKitterick, Fatema-tuz Johura, Marzia Sultana, Tahmeed Ahmed, Munirul Alam, and Kimberley D Seed. Temporal shifts in antibiotic resistance elements govern phage-pathogen conflicts. Science, 373(6554):eabg2166, 2021.

28. Rachel AF Wozniak and Matthew K Waldor. Integrative and conjugative elements: mosaic mobile genetic elements enabling dynamic lateral gene flow. Nature Reviews Microbiology, 8(8):552–563, 2010.

29. Julie D Thompson, Toby J Gibson, and Des G Higgins. Multiple sequence alignment using clustalw and clustalx. Current protocols in bioinformatics, (1):2–3, 2003.

30. Kaitlin Chaung, Tavor Z Baharav, George Henderson, Peter Wang, Ivan N Zheludev, and Julia Salzman. A statistical reference-free algorithm subsumes and generalizes common genomic sequence analysis and uncovers novel biological regulation. bioRxiv, pages 2022–06, 2022.

31. Yoav Benjamini and Daniel Yekutieli. The control of the false discovery rate in multiple testing under dependency. Annals of statistics, pages 1165–1188, 2001.

32. Christiam Camacho, George Coulouris, Vahram Avagyan, Ning Ma, Jason Papadopoulos, Kevin Bealer, and Thomas L Madden. Blast+: architecture and applications. BMC bioinformatics, 10(1):1–9, 2009.

33. Jessica Storer, Robert Hubley, Jeb Rosen, Travis J Wheeler, and Arian F Smit. The dfam community resource of transposable element families, sequence models, and genome annotations. Mobile DNA, 12(1):1–14, 2021.

34. Raphael Leplae, Gipsi Lima-Mendez, and Ariane Toussaint. Aclame: a classification of mobile genetic elements, update 2010. Nucleic acids research, 38(suppl_1):D57–D61, 2010.

35. Monica Santamaria, Bruno Fosso, Flavio Licciulli, Bachir Balech, Ilaria Larini, Giorgio Grillo, Giorgio De Caro, Sabino Liuni, and Graziano Pesole. Itsonedb: a comprehensive collection of eukaryotic ribosomal rna internal transcribed spacer 1 (its1) sequences. Nucleic Acids Research, 46(D1): D127–D132, 2018.

36. Christian Selig, Matthias Wolf, Tobias Müller, Thomas Dandekar, and Jörg Schultz. The its2 database ii: homology modelling rna structure for molecular systematics. Nucleic acids research, 36(suppl_1):D377–D380, 2007.

37. Meng Liu, Xiaobin Li, Yingzhou Xie, Dexi Bi, Jingyong Sun, Jun Li, Cui Tai, Zixin Deng, and Hong-Yu Ou. Iceberg 2.0: an updated database of bacterial integrative and conjugative elements. Nucleic acids research, 47(D1):D660–D665, 2019.

38. Ioanna Kalvari, Eric P Nawrocki, Nancy Ontiveros-Palacios, Joanna Argasinska, Kevin Lamkiewicz, Manja Marz, Sam Griffiths-Jones, Claire Toffano-Nioche, Daniel Gautheret, Zasha Weinberg, et al. Rfam 14: expanded coverage of metagenomic, viral and microrna families. Nucleic Acids Research, 49(D1):D192–D200, 2021.

