## Supplemental Information for "DIVE: a reference-free statistical approach to diversity-generating & mobile genetic element discovery"

#### 1 Algorithmic details

DIVE operates symmetrically, forming upstream and downstream scores for each anchor; we describe the procedure here for downstream targets. For each anchor, DIVE generates a target dictionary with an online clustering method that collapses targets within "sequencing error" distance. It then models the number of clusters formed at each step using a Poisson-Binomial model. An anchor must be observed sufficiently often ($> N_{\min}$ times) to have adequate statistical power to call its targets more diverse than expected by chance. To keep the memory footprint low, for every 100k reads, DIVE computes the probability that an anchor will be observed at least $N_{\min}$ times in the FASTQ file ($N_{\min} = 25$). If this probability is $< 1\%$, DIVE does not accept newly observed anchors into the dictionary. After the FASTQ file is completely traversed, DIVE uses a Poisson-Binomial model to compute the probability $P_{H_0}(C_N \geq c_N \mid N)$ that the number of clusters $C_N$ exceeds the observed value $c_N$ under the null hypothesis that the observed target diversity is due to background variability. DIVE uses the Benjamini-Hochberg (BH) correction to control the false discovery rate (FDR). Anchors producing significant target diversity are clustered using DBSCAN with the Levenshtein edit distance as a metric to remove redundancies. Within each anchor cluster, the anchor with the largest effect size in each direction is picked as the *representative anchor*. The analysis in this paper has been done using the representative anchors. Representative anchors, the number of clusters ($C$), the number of times a target was observed with the anchor ($N$), the effect size $\alpha(c_N) = \log_2(c_N / E[C_N \mid N])$, and corrected $p$-values are reported. All numbers described above, such as k, 1%, and 100k, are user-adjustable parameters. To facilitate interpretation, DIVE aligns the RAs (BLAST) to a set of databases provided by the user.

#### 2 Removal of technical artifacts

Artificial sequences such as sequencing adapters will exhibit statistical signatures like bona fide MGEs (highly variable downstream sequences) and serve as positive controls. DIVE identified several such sequences, and we removed them before proceeding to the rest of the downstream analysis. We also discarded representative anchors with low nucleotide entropy ($H < 3$) to remove other potential technical artifacts. To that end, we compute an empirical estimate of the entropy based on 5-mers using

$$H = \sum_{x \in \mathcal{S}_5} p(x) \log p(x)$$

where *p(x)* is estimated from the anchor by partitioning it into overlapping 5-mers and where we take the convention that $0 \log 0 = 0$. We require $H > 3$ for the anchor to be considered in the downstream analysis.

### 3 Simulations

In order to test the performance of DIVE in detecting the termini of mobile genetic elements, we performed two sets of simulations (Fig. S2).

In the first setting, we simulated an ancestral mobile element inserted in the same position in all the sampled genomes (Fig. S2a). First, a base random genome of length 100 kbp is generated. Then, mutations and indels are randomly introduced to generate 100 different genomes, modeling the strain variation of the organism. This is done for varying rates MIR$= 0.01, 0.025, 0.05, 0.1, 0.2$ at each nucleotide in each individual genome. If the original genome is modified, then with probability 0.8 a substitution is introduced and with probability 0.2 an insertion or deletion is introduced with equal probability. This results in a set of 100 genomes with varying degrees of sequence divergence. Subsequently, a fixed number of copies of a randomly generated sequence of 100 bp, representing an ancestral mobile element, is introduced into each individual genome with varying copy number CN$= 2, 5, 10, 15, 20$. Finally, the individual genomes are sampled to obtain 100 bp reads producing the desire coverage C$= 5, 10, 20, 30, 40, 50$. This results in a set of reads that are used as input for DIVE. The anchors identified by DIVE are then compared to the termini of the simulated mobile element to determine the specificity and sensitivity of the algorithm. A true positive consists of a sequence overlap greater than 10 nt. This procedure is done 250 times for each combination of mutation/indel rate MIR, copy number CN, and coverage C, which allows us to calculate the true positive rate and the false positive rate for a given $q$-value threshold. Varying the $q$ threshold we obtained the ROC shown in Fig. S3.

In the second setting, we simulated an active mobile element with a single copy inserted in different positions among the sampled genomes (Fig. S2b). We follow the same procedure as in the first scenario to generate a set of 100 genomes with varying degrees of sequence divergence (MIR$= 0.01, 0.025, 0.05, 0.1, 0.2$). Subsequently, a single copy of a randomly generated sequence of 100 bp, representing an active mobile element, is introduced into each individual genome. A new position is selected for each genome with varying probabilities, emulating different levels of activity of the element (AL$= 0.25, 0.5, 0.75, 1.0$). Finally, the individual genomes are sampled to obtain 100 bp reads producing the desire coverage C$= 5, 10, 20, 30, 40, 50$. This results in a set of reads that are used as input for DIVE. The anchors identified by DIVE are then compared to the termini of the simulated mobile element to determine the specificity and sensitivity of the algorithm. A true positive consists of a sequence overlap greater than 10 nt. This procedure is done 250 times for each combination of mutation/indel rate MIR, activity level AL, and coverage C, which allows us to calculate the true positive rate and the false positive rate for a given $q$-value threshold. Varying the $q$ threshold we obtained the ROC shown in Fig. S4.

Note that in both cases the performance of DIVE improves with the coverage as expected. In the ancestral mobile element simulation, the performance also improves with higher copy number, whereas in the active element simulation the performance there is an improvement for higher activity levels.

### 4 Comparison with MGEfinder

MGEfinder [1] is a recently developed tool for *de novo* discovery of MGEs that outperformed the state of the art methods for mobile element discovery. As the rest of the tools, however, MGEfinder requires a reference genome to perform an alignment step. As a result, the quality of the reference severely conditions the algorithm's sensitivity. In parallel, MGEfinder relies on assembly algorithms to assemble all the available sequencing reads. To detect MGEs, the algorithm searches for clipped aligned reads to identify putative termini of mobile elements. Since this is the critical part of the algorithm, we compared the performance of both DIVE and MGEfinder in the previous simulated scenarios.

We first used the previous simulated data to compare the performance of the two tools in detecting the termini of mobile elements, emulating the simulations performed in [1]. To that end, we ran MGEfinder (v1.0.6) on each set of reads generated during our simulations. Note, however, that MGEfinder does not compute a p-value, and thus we cannot produce ROC curves and compare the area under the curve of the two algorithms. Instead, we limited the comparison to the true positive rate of the algorithms, using a significance threshold of $q \leq 0.1$ for DIVE, which we also use in the rest of the paper. As for MGEfinder, the reference genome used in the one generated in the first step (see Fig. S2), and the parameters were just set to the default values.

In the first scenario where we considered an ancient mobile element with multiple copies in the genome, MGEfinder only outperfomed DIVE when there are only two copies of the element and when the mutation and indel rate was very low. In the rest of scenarios, DIVE performed at least as well as MGEfinder. When the mutation and indel rate is greater than 1%, DIVE consistently shows higher sensitivity than MGEfinder, which eventually cannot detect the mobile element (TPR=0%) due to the lack of reliability of the reference genome. In contrast, the performance of DIVE is not altered by the mutation and indel rate since it does not rely on an alignment step to a particular reference genome. In the second scenario, where the mobile element has a single copy found at different loci at different rates, DIVE can detect the termini with sufficient coverage (20x), whereas MGEfinder cannot do so consistently even at low mutation and indel rates. Overall, DIVE shows greater sensitivity than MGEfinder in almost all scenarios in detecting mobile element termini, which is the foundation of both algorithms.

Nevertheless, to test if the increased theoretical power of DIVE compared to MGEfinder is born out in real data, we ran DIVE on data highlighted in the MGEfinder study. We analyzed 200 *Neisseria gonorrhoeae* isolates used in [1] and compared the detection power of DIVE to reported results by MGEfinder. DIVE reported 81,792 unique RAs mapping to 190 different TEs (BLAST $e < 0.01$), compared to 28 TEs reported by MGEfinder in the 891 isolates analyzed. TEs detected by DIVE could constitute ancestral TEs present in the *N. gonorrhoeae* genome. To test this, we ran DIVE on unaligned reads after mapping (bowtie2) to the reference genome as in [1] (overall alignment rate 88%). DIVE called 868 unique RAs mapping to 107 different TEs (Table S4, Fig. 2). The large effect sizes found in reads failing to align to the reference genome lead us to hypothesize that these elements are active TEs in *N. gonorrhoeae*. Unfortunately, however, the identity of the TEs detected by DIVE and MGEfinder were not comparable since the latter did not cross-reference their predicted TEs with any database.

### 5 Novel CRISPR array in *R. bromii*

We ran DIVE on human gut metagenomic samples with a *Rotavirus* infection (Table S13). DIVE called 133,528 unique RAs mapping to 2,991 MGEs and 575 CRISPR repeats. Imposing the CRISPR repeat criteria outlined in the main text revealed an RA that did not perfectly match any known repeat in the CRISPR-Cas++ repeat database but aligned 15 times (BLAST $e < 0.01$) to a phage identified in metagenomic data [2]. The RA was not found in any other sequence on the NCBI nucleotide collection (nr/nt) or the DR database but has 2 mismatches from the closest DR. We note that BLAST of the RA to the NCBI WGS database restricted to the bacterium *Ruminococcus bromii* (taxid:40518) gives perfect matches in a locus with CRISPR-associated (Cas) genes very similar to that in the phage contig (other parts of the phage contig also match to *R. bromii*). 13/43 targets mapped (BLAST $e < 0.01$) to *Siphoviridae* (13/13), *Podoviridae*(2/13), and *Myoviridae* (1/13), supporting it as a CRISPR system. Thus, we hypothesize this RA constitutes a novel functional CRISPR DR. To the best of our knowledge, no CRISPR system has been previously reported in *R. bromii*.

### 6 Non-coding RNA as magnets of MGEs

In *Escherichia coli*, DIVE called 712 unique RAs that showed target hyper-variability mapping to 84 different Rfam accessions annotated as tRNA genes (BLAST $e < 0.01$). The target diversity of seven RAs, mapping to four unique tRNA genes, was significantly associated with the isolation source (Table S7; Fig. S5). These included Met-tRNA, a known target of retrotransposons [3], and a valU operon encoding three tRNA genes. The RAs tended to align to the ends of the gene and accrued up to 3,000 unique targets resulting in 1,400 clusters (Methods). However, no RA mapped within an individual genome on the NCBI nucleotide database more than eleven times (BLAST $e < 0.01$), suggesting the observed target variability cannot be explained alone by the within-genome variability. Only 2% of the targets mapped tRNA genes (bowtie2), suggesting the observed variability was not due to subtle modifications in the tRNA sequence.

Among ncRNA genes showing a significant association with the isolation source, we identified an *E. coli* nucleoid-associated noncoding RNA 4 (naRNA4) gene (U00096.3/900752-900832). naRNA4 genes are encoded in repetitive extragenic palindrome (REP) regions, previously observed at the recombination junctions of lambda phages [4] and described as hotspots for transposition events [5]. 823 unique RAs mapped (BLAST $e < 0.01$) to 82 Rfam accessions annotated as distinct naRNA4 genes, concentrating at the ends of the gene (Table S8). The maximum number of times a RA aligned within an individual genome on NCBI was 131 (BLAST $e < 0.01$), whereas RAs had up to 140,000 distinct targets, comprising over 500 sequence clusters. Furthermore, over 90% of targets did not align to any naRNA4 gene (bowtie2), suggesting the observed variability was not due to subtle modifications to the naRNA4 sequence. DIVE produced similar results for four tRNA genes in the *Vibrio cholera* isolate data (Methods). DIVE also identified 264 RAs mapping (BLAST $e < 0.01$) to 26 Rfam accessions annotated as antisense RNA 5 (asRNA5), including four RAs mapping to two *V. cholerae* O1 biovar el Tor asRNA5 genes showing a target diversity significantly associated with available covariates (Table S9). RAs showed no bias in their alignment position. Although no RA mapped within an individual genome on NCBI more than 51 times (BLAST $e < 0.01$), some RAs reached 160 target clusters. Furthermore, over 75% of targets did not map to any asRNA5 gene (bowtie2), suggesting the observed target diversity was not due to subtle modifications to the asRNA5 sequence.

### 7 Antibiotic resistance in *V. cholerae* SXT ICEs

We ran DIVE on 247 *V. cholerae* isolates and aligned the resulting RAs to six known SXT ICEs. RAs accumulated nearby antibiotic resistance genes (Fig. 2), which define cargo gene hotspots. In particular, RAs clustered near known hotspots in SXT ICE variants ICE*Vch*Ind4, ICE*Vch*Ind5, ICE*Vch*MO10, ICE*Vch*Ban9, and ICE*Vch*Chn956. In addition, we found a unique RA cluster in ICE*Vch*Ban8 in a different location, overlapping a known transposase. Furthermore, when we aligned the corresponding set of targets to a TE database (TnCentral) [6], we found alignment rates of 10% (bowtie2), consistent with the fact that cargo genes mobilize via transposons [7]. Only targets derived from RAs aligning to ICE*Vch*Ban8, a variant lacking annotation information on the database, produced an alignment rate of 0.02%. Nevertheless, in 19% of the targets of RAs aligning to this SXT variant, we observed $[A]^n$ or $[T]^n$ homopolymers, with $n \geq 3$, at the ends of the targets, consistent with the insertion signature of non-LTR transposons. Thus, we hypothesize these RAs point to a putative hotspot in ICE*Vch*Ban8. The percentage of targets among the other SXT variants enriched in A/T homopolymers ranged from 25 to 31%.

### 8 Unexplained genetic diversity in *E. coli* and *V. cholerae*

In *E. coli*, RA CCGCCATATCACCTCCGTGATGGTTGC showed the largest median effect ($\alpha = 4.22$, prevalence=5.28%). The RA produced a single match in at least one hundred NCBI accessions, mainly of *E. coli* strains. In at least three references, the RA lies within the coding sequence of the KdsD gene (arabinose-5-phosphate isomerase), approximately 60 bp from the stop codon (Fig. 3). Only two targets were observed upstream (n=514), but 1,653 different targets (697 clusters) were observed downstream. We found strong agreement upstream of the RA across reads (¿99% nucleotide identity) and a significant decay downstream in nucleotide identity across reads, with some positions showing less than 35% agreement (Fig. 3). This diversity was not well represented on the reference genomes on NCBI, with only 21 targets (1% of different target sequences) producing hits on the database (BLAST $e < 0.5$). In addition, sequence hyper-variability extended well beyond the range of the target sequence, spanning over

100 nt, and the variation included substitutions and indels altering the protein sequence of KdsD.

In *V. cholerae*, RA CGACTGGTTAAACCACAACCAAATGCA showed the largest median effect size ($\alpha = 3.47$, prevalence=5.17%). The RA produced a single match in at least one hundred NCBI accessions, all *V. cholerae* strains, mapping to the intergenic region between tRNA-Cys and the artP gene (arginine ABC transporter ATPase), with the former being in the direction of more target diversity (Fig. 3). The RA only had two upstream targets but 396 different targets (95 clusters) downstream. We found strong agreement upstream of the RA across reads (¿99% nucleotide identity) and a significant decay downstream in nucleotide identity across reads, with some positions showing less than 45% agreement (Fig. 3). This diversity was not well represented on the reference genomes on NCBI, with only 61 targets (15% of different target sequences) producing hits on the database (BLAST $e < 0.5$). In addition, sequence hyper-variability extended well beyond the range of the target sequence, spanning over 100 nt, and the variation included substitutions and indels (Fig. 3).

### References


[1] M. G. Durrant, M. M. Li, B. A. Siranosian, S. B. Montgomery, and A. S. Bhatt, "A bioinformatic analysis of integrative mobile genetic elements highlights their role in bacterial adaptation," *Cell host & microbe*, vol. 27, no. 1, pp. 140–153, 2020.

[2] M. J. Tisza and C. B. Buck, "A catalog of tens of thousands of viruses from human metagenomes reveals hidden associations with chronic diseases," *Proceedings of the National Academy of Sciences*, vol. 118, no. 23, p. e2023202118, 2021.

[3] G. Martinez, "trnas as primers and inhibitors of retrotransposons," *Mobile Genetic Elements*, vol. 7, no. 5, pp. 1–6, 2017.

[4] M. Kumagai and H. Ikeda, "Molecular analysis of the recombination junctions of $\lambda$ bio transducing phases," *Molecular and General Genetics MGG*, vol. 230, no. 1, pp. 60–64, 1991.

[5] R. Tobes and E. Pareja, "Bacterial repetitive extragenic palindromic sequences are dna targets for insertion sequence elements," *BMC genomics*, vol. 7, no. 1, pp. 1–12, 2006.

[6] K. Ross, A. M. Varani, E. Snesrud, H. Huang, D. O. Alvarenga, J. Zhang, C. Wu, P. McGann, and M. Chandler, "Tncentral: a prokaryotic transposable element database and web portal for transposon analysis," *MBio*, vol. 12, no. 5, pp. e02060–21, 2021.

[7] S. Benler, G. Faure, H. Altae-Tran, S. Shmakov, F. Zhang, and E. Koonin, "Cargo genes of tn 7-like transposons comprise an enormous diversity of defense systems, mobile genetic elements, and antibiotic resistance genes," *Mbio*, vol. 12, no. 6, pp. e02938–21, 2021.

### Supplementary Figures

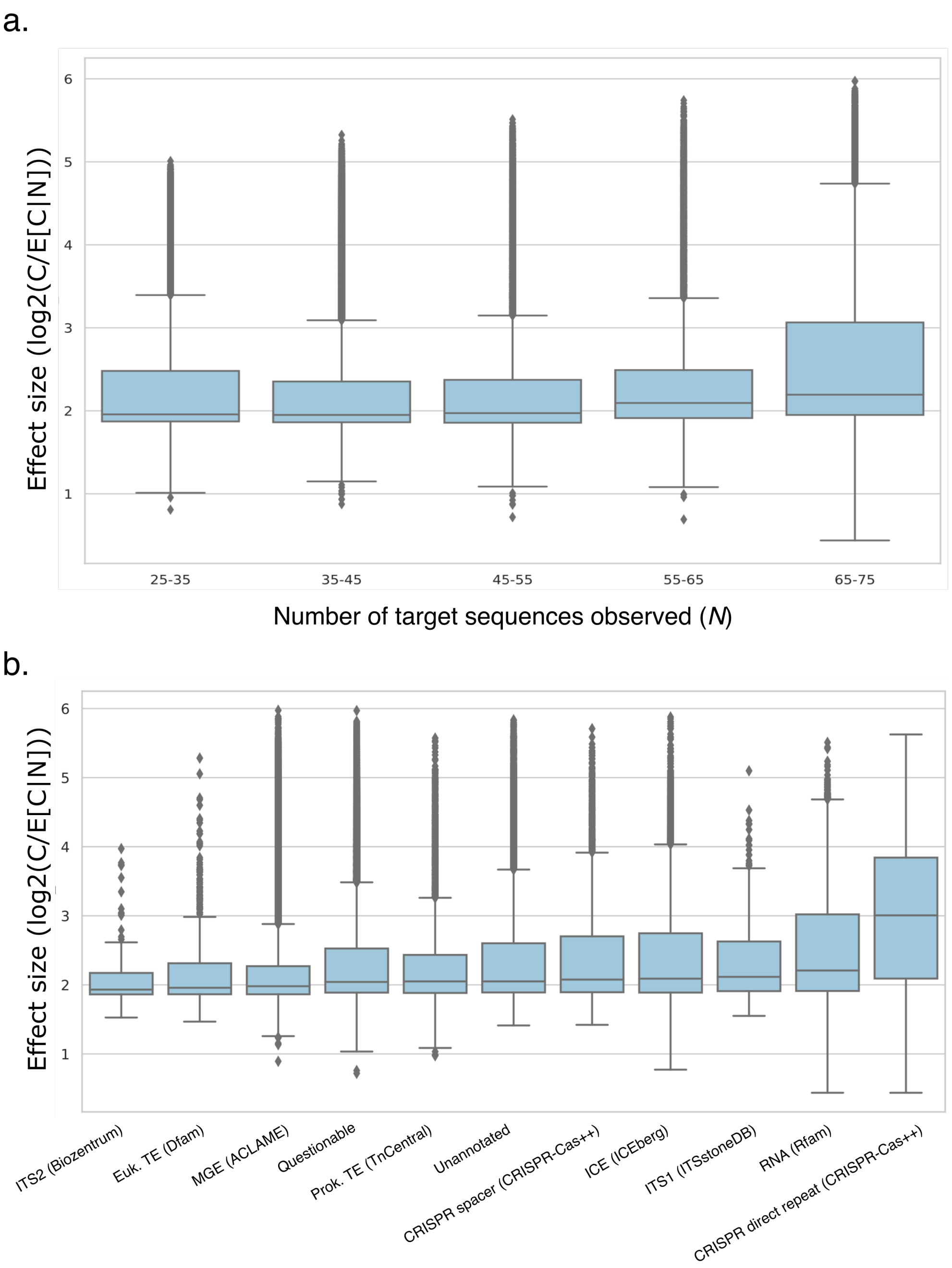


**Figure S1: DIVE effect size. a.** Boxplot of the effect size $\log_2(c_N/E[C|N])$ computed by DIVE as a function of the number of target sequences observed $N$ in *E. coli* (binned in intervals of 10). We designed the effect size to remove any dependence on $N$, as would happen if we simply looked at the number of clusters $C$. As the plot shows, the effect size is not confounded by the number of target sequences observed $N$ (OLS $R^2 = 0.02$). **b.** Boxplot of the median effect size (across anchors) of the elements within each category, with anchors mapping to known CRISPR direct repeats showing the highest median effect size.

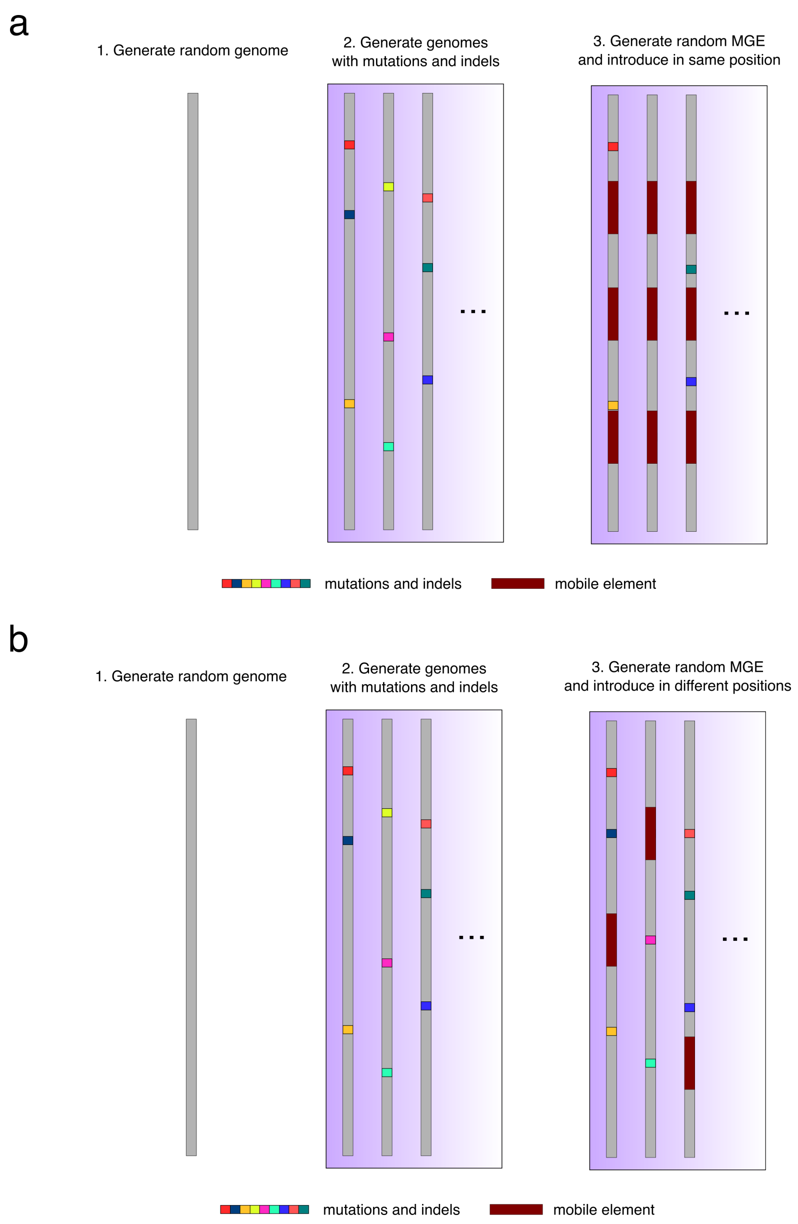


**Figure S2: Ancestral and active mobile element simulations a.** An ancestral mobile element is introduced at the same position in each genome. A random genome of length 100 kbp is generated first. Then, mutations and indels are randomly introduced to generate 100 different genomes. A fixed number of copies of an ancestral mobile element is introduced into each one of the simulated genomes. Finally, the genomes are sampled to produce 100 bp reads. The rate of mutations and indels, the reads produced, and the number of copies are varied throughout the simulations. **b.** An active mobile element is introduced in different positions in each genome. A random genome of length 100 kbp is generated first. Then, mutations and indels are randomly introduced to generate 100 different genomes. A single copy of the element is introduced in a new position in each genome with some probability. Finally, the genomes are sampled to produce 100 bp reads. The rate of mutations and indels, the number of reads produced, and the probability of the insertion in a new locus are varied throughout the simulations.

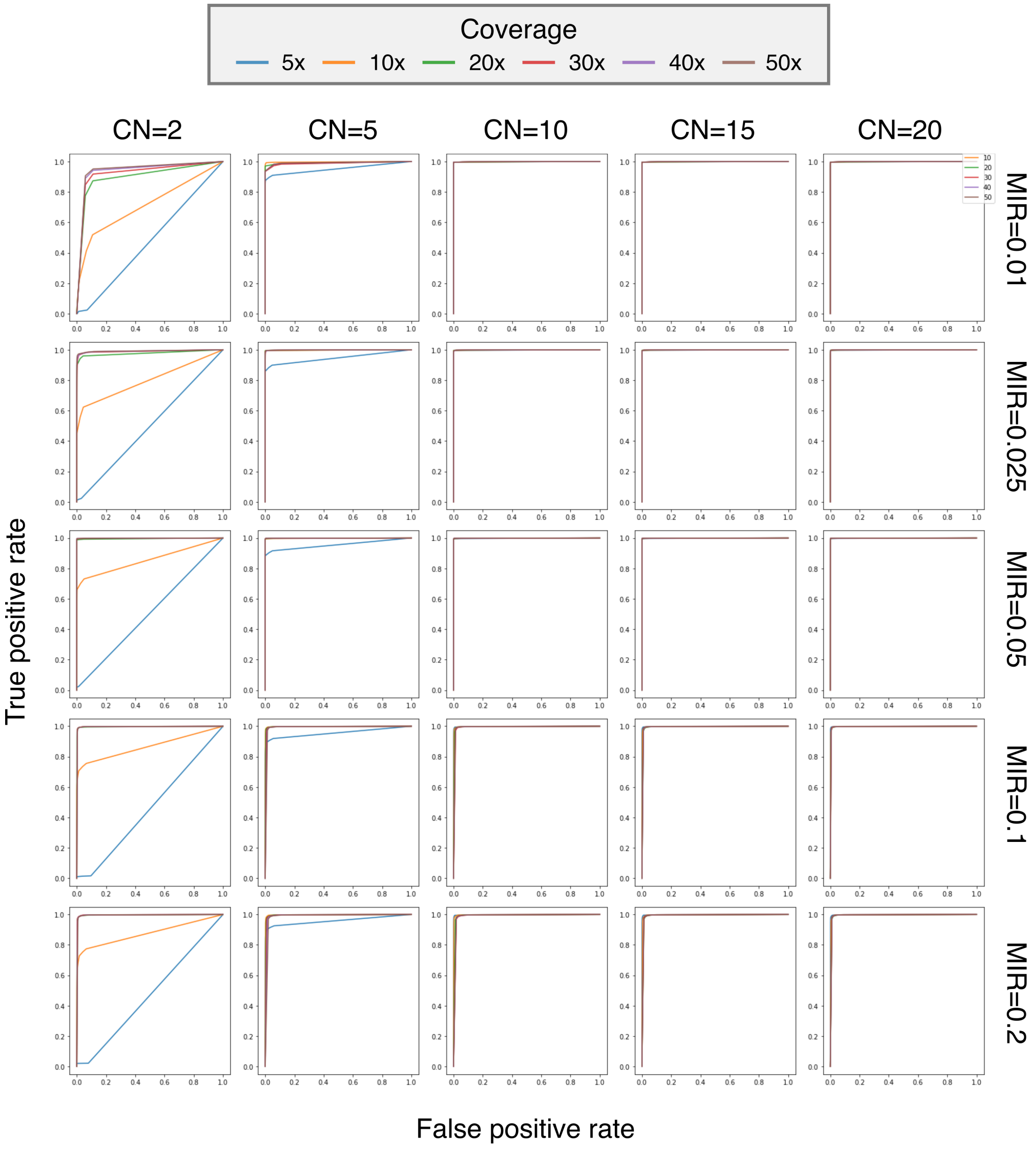


**Figure S3: DIVE ROC in ancestral mobile element simulations.** Receiving operating curve (ROC) of DIVE in detecting the termini of an ancestral mobile genetic element with varying coverage levels, copy numbers (CN) and varying degrees of genome divergence as quantified by the mutation and indel rate (MIR).

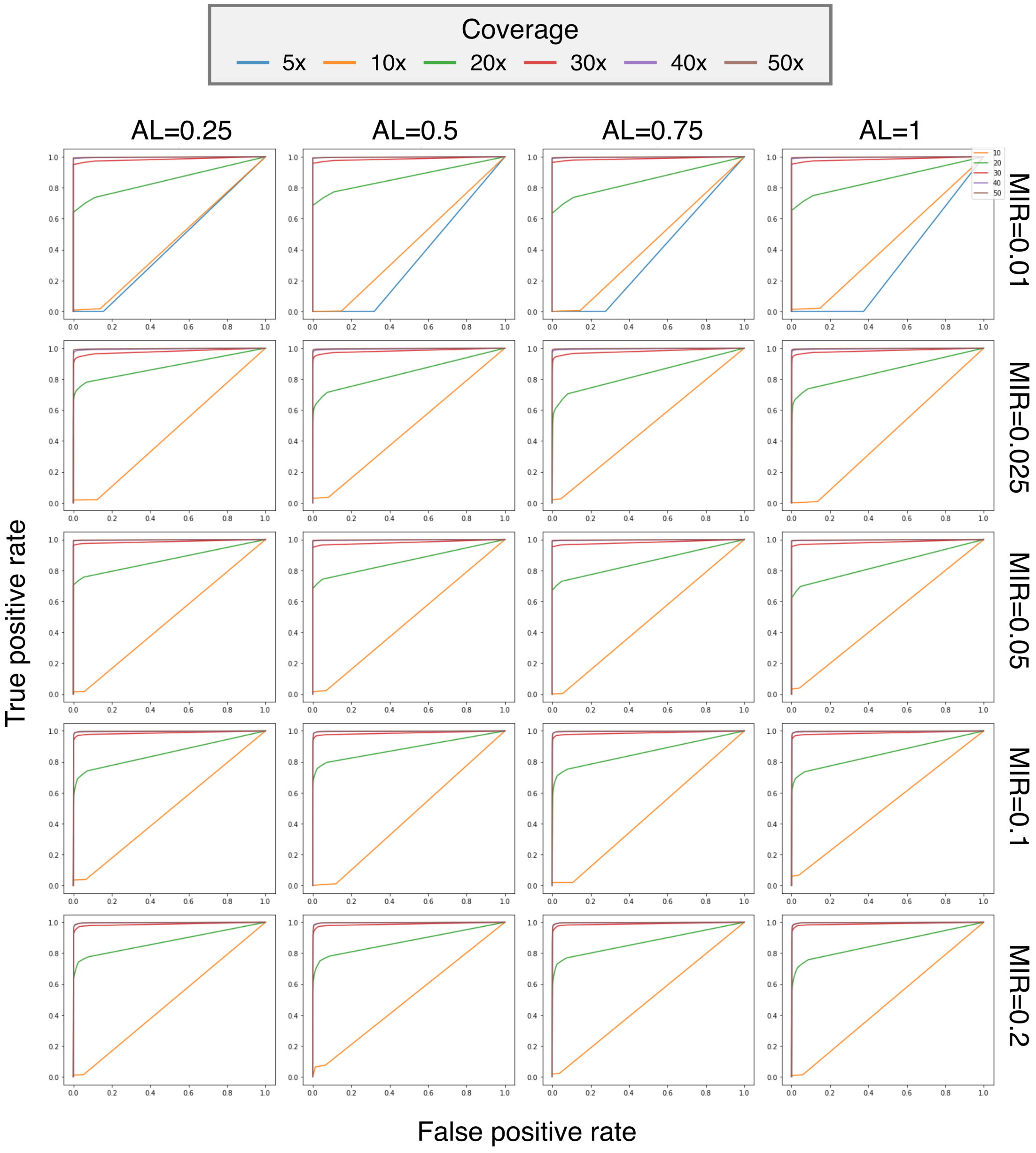


**Figure S4: DIVE ROC in active mobile element simulations.** Receiving operating curve (ROC) of DIVE in detecting the termini of an active mobile genetic element with varying coverage levels, activity levels (AL) and varying degrees of genome divergence as quantified by the mutation and indel rate (MIR).

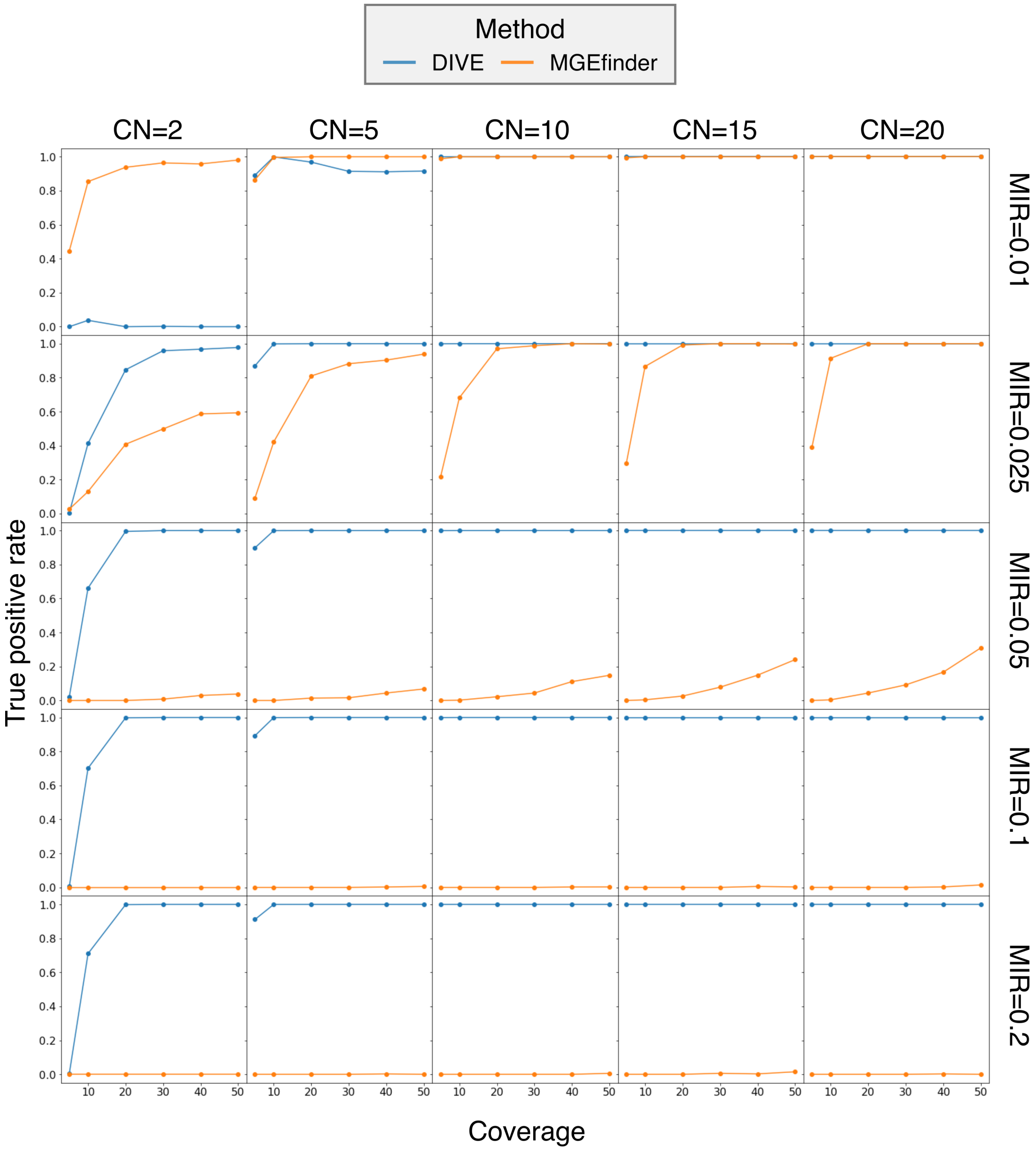


**Figure S5: True positive rate DIVE vs MGEfinder (ancestral element).** True positive rate (sensitivity) of DIVE (blue) and MGEfinder (orange) in detecting the termini of an ancestral mobile genetic element with varying coverage levels, copy numbers (CN) and varying degrees of genome divergence as quantified by the mutation and indel rate (MIR).

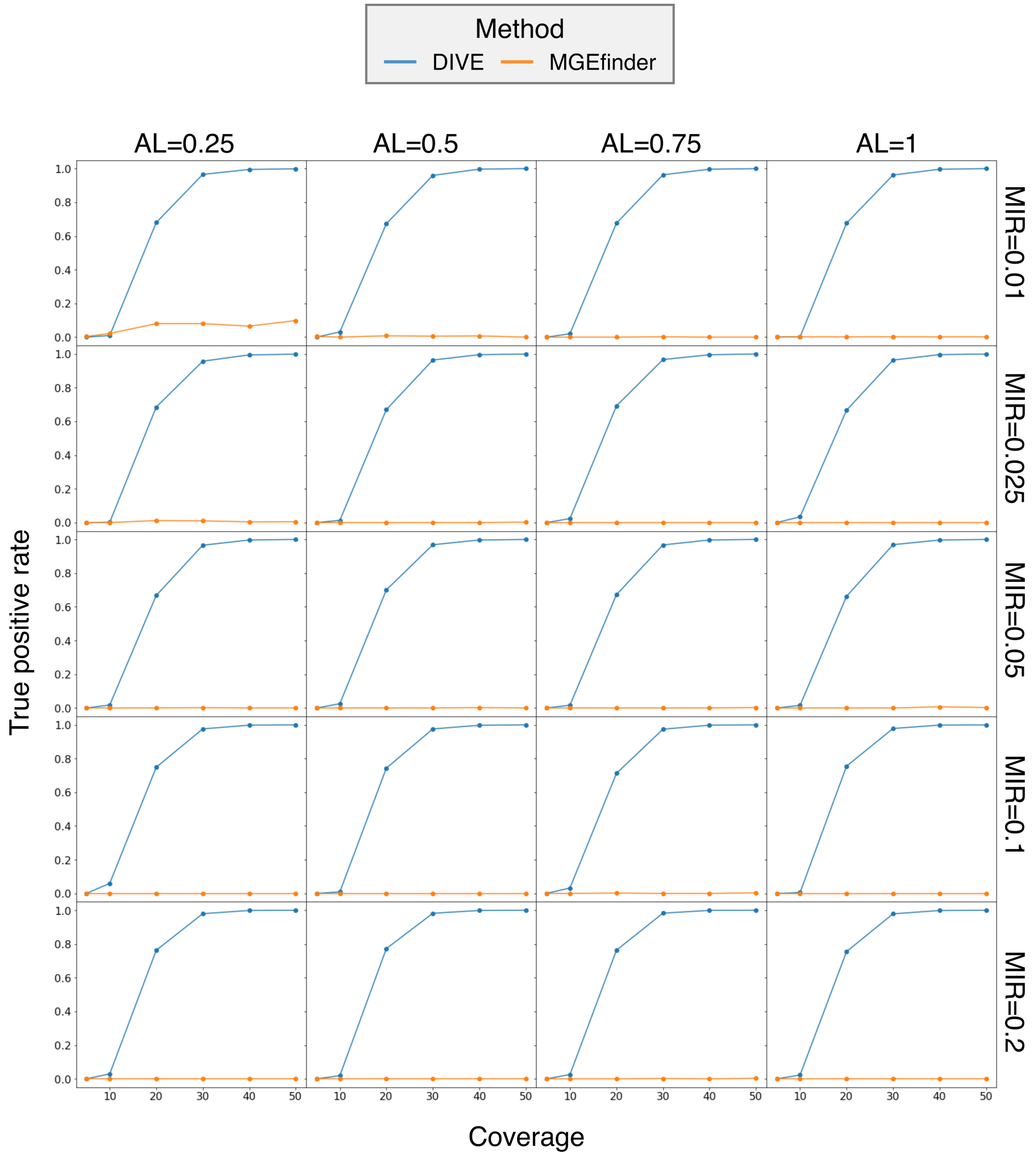


**Figure S6: True positive rate DIVE vs MGEfinder (active element).** True positive rate (sensitivity) of DIVE (blue) and MGEfinder (orange) in detecting the termini of an active mobile genetic element with varying coverage levels, activity levels (AL) and varying degrees of genome divergence as quantified by the mutation and indel rate (MIR).

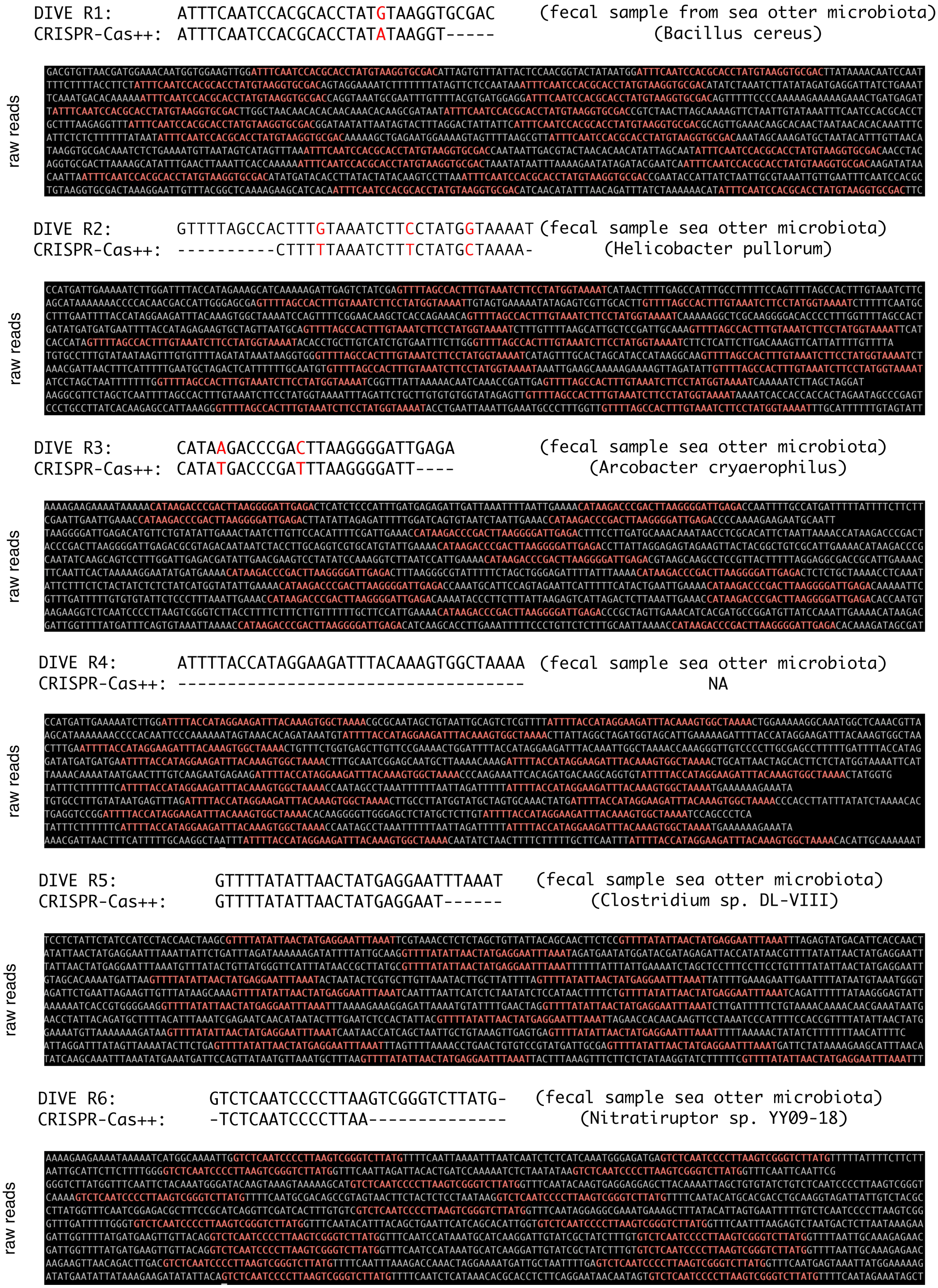


**Figure S7:** ***De novo* discovery of CRISPR repeats**. Each row shows the putative CRISPR repeat identified from DIVE's output (DIVE R), the closest match in the CRISPR-Cas++ database when found, and a subset of reads showing the repetition of the repeat within single reads.

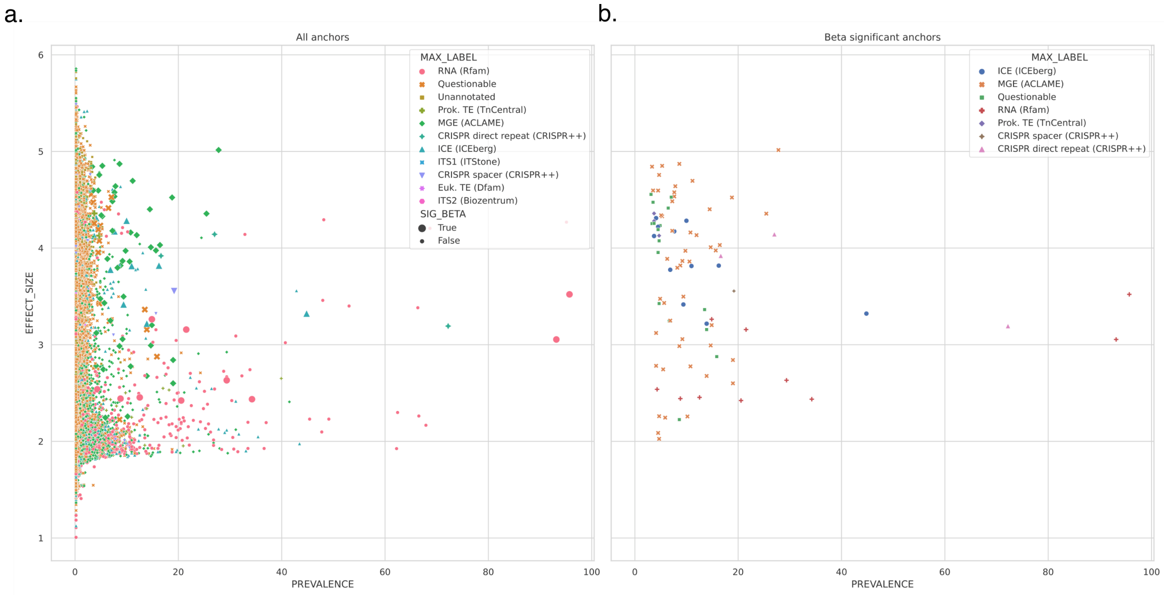


**Figure S8: Median effect size of each element and its prevalence across the *E. coli* dataset. a.** Median effect size of each element (mapped by at least an anchor flagged by DIVE) as a function of its prevalence across samples. Elements mapped by an anchor with target diversity significantly associated with the available covariates are shown larger. **b.** Median effect size of each element as a function of its prevalence across samples for those cases where the target diversity of an anchor was significantly associated with the available covariates.

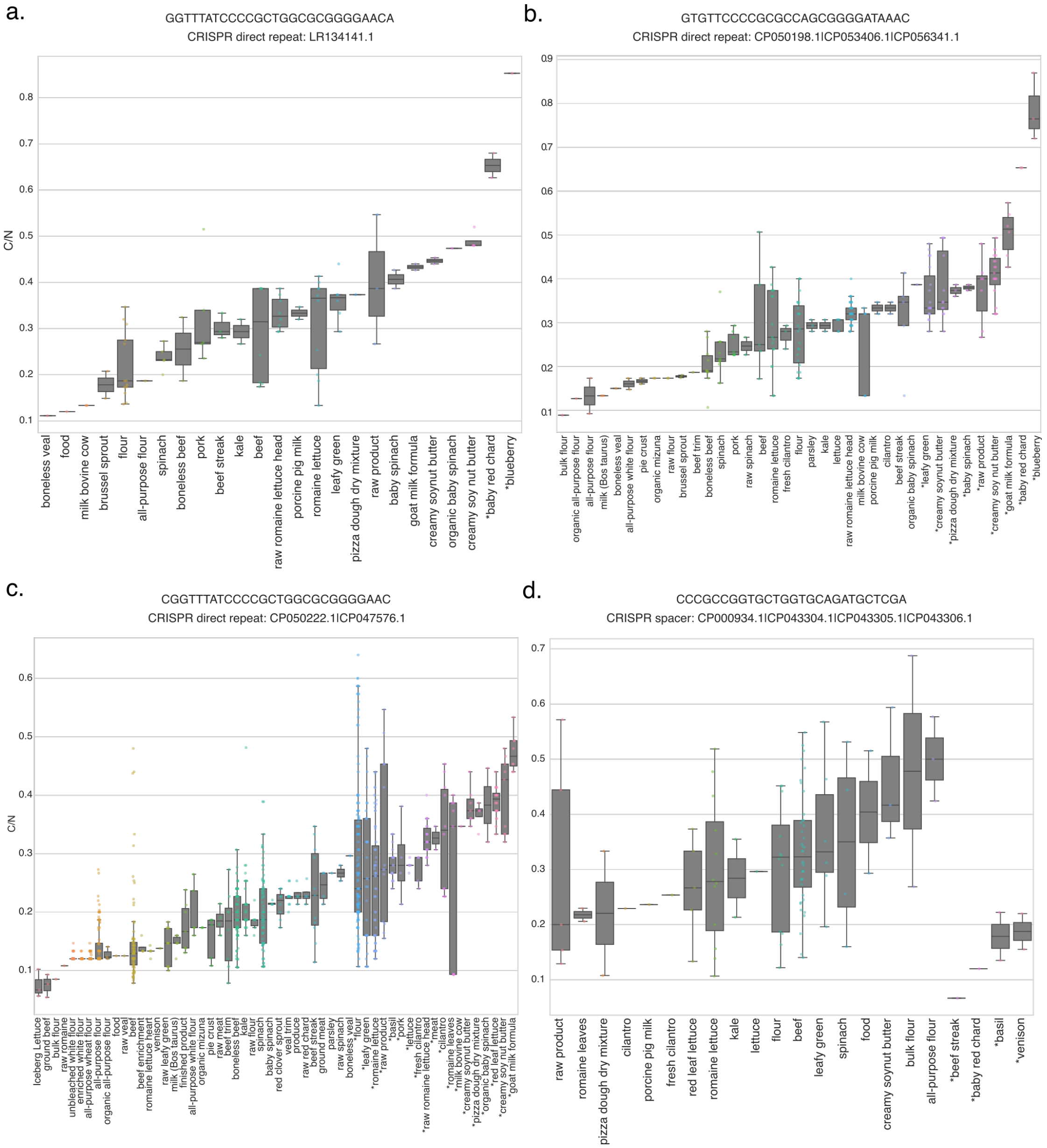


**Figure S9: Spacer turnover in CRISPR arrays is a function of the isolation source in *E. coli* isolates.** **a.** Anchor GGTTTATCCCCGCTG-GCGCGGGGAACA, mapping to CRISPR direct repeat LR134141.1, shows an almost 10-fold difference in target diversity between isolates obtained from blueberry than that of veal. **b.** Anchor GTGTTCCCCGCGCCAGCGGGGATAAAC, mapping to CRISPR direct repeat CP050198.1—CP053406.1—CP056341.1 shows an almost 10-fold difference in target diversity between isolates obtained from blueberry than that of flour. **c.** Anchor CGGTTTATCCCCGCTGGCGCGGGGAAC, mapping to CRISPR direct repeat CP050222.1—CP047576.1 shows an almost 10-fold difference in target diversity between isolates obtained from goat milk formula than that of iceberg lettuce. **d.** Anchor CCCGCCGGTGCTGGTGCAGATGCTCGA, mapping to CRISPR spacer CP000934.1—CP043304.1—CP043305.1—CP043306.1 shows a 5-fold difference in target diversity between isolates obtained from all-purpose flour than that of beef steak.

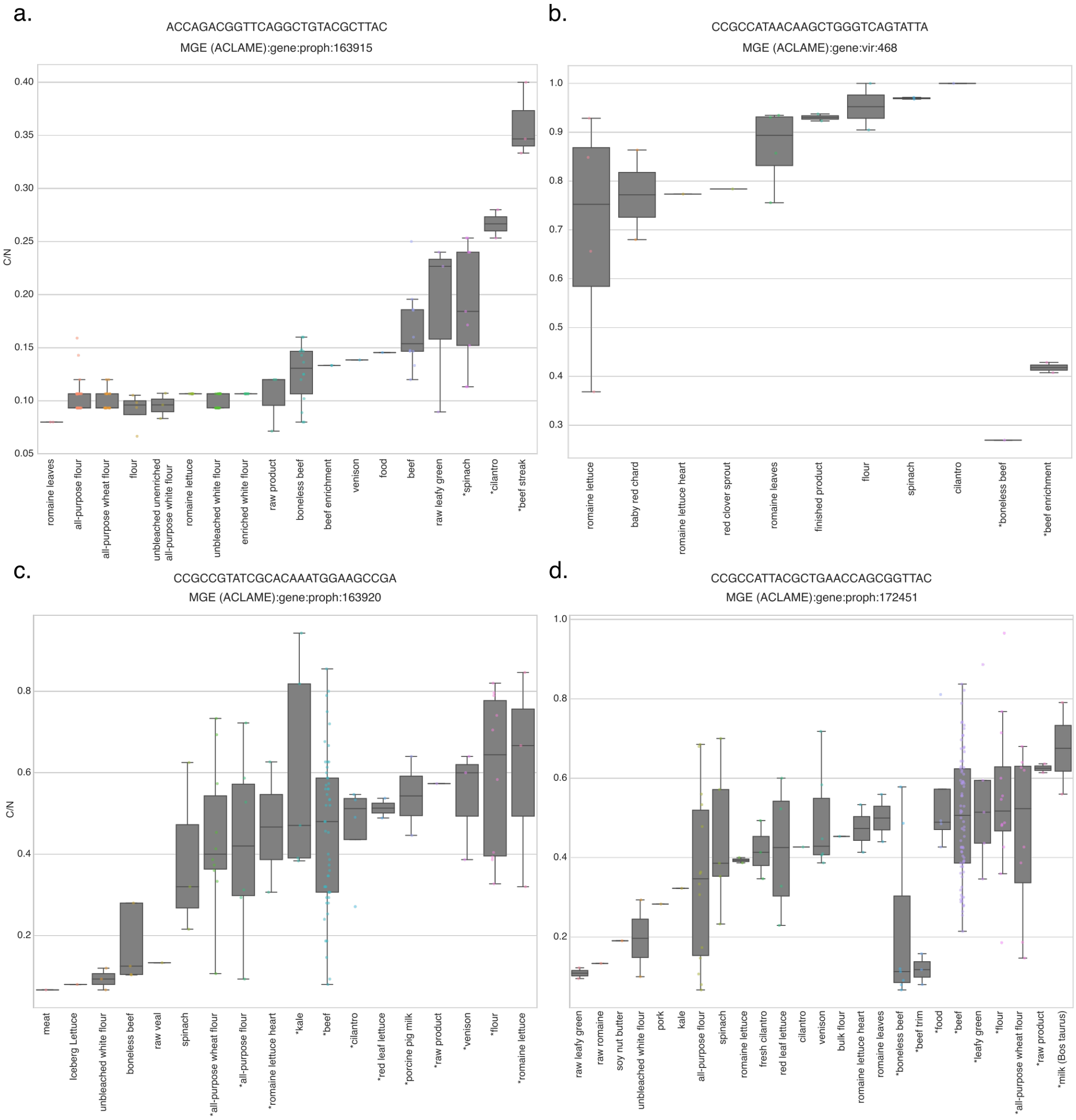


**Figure S10: Mobile genetic elements showing different mobility levels as a function of the isolation source in *E. coli* isolates. a.** Anchor ACCAGACGGTTCAGGCTGTACGCTTAC, mapping to a known MGE (ACLAME: 163915), shows a 5-fold difference in target sequence diversity between isolates obtained from beef steak and that of romaine lettuce. Upon BLAST search, we determined this element to be an IS66-family insertion sequence. **b.** Anchor CCGCCATAACAAGCTGGGTCAGTATTA, mapping to a known MGE (ACLAME: 468), shows a 4-fold difference in target sequence diversity between isolates obtained from boneless beef and that of cilantro. Upon BLAST search, we determined this element to be a lambdoid phage. **c.** Anchor CCGCCGTATCGCACAAATGGAAGCCGA, mapping to a known MGE (ACLAME: 163920), shows a 6-fold difference in target sequence diversity between isolates obtained from meat and that of romaine lettuce. Upon BLAST search, we determined this element to be a IS66-family insertion sequence. **d.** Anchor CCGCCATTACGCTGAACCAGCGGTTAC, mapping to a known MGE (ACLAME: 172451), shows a 6-fold difference in target sequence diversity between isolates obtained from raw leafy green and that of *Bos taurus* milk. Upon BLAST search, we determined this element to be a IS66-family insertion sequence.

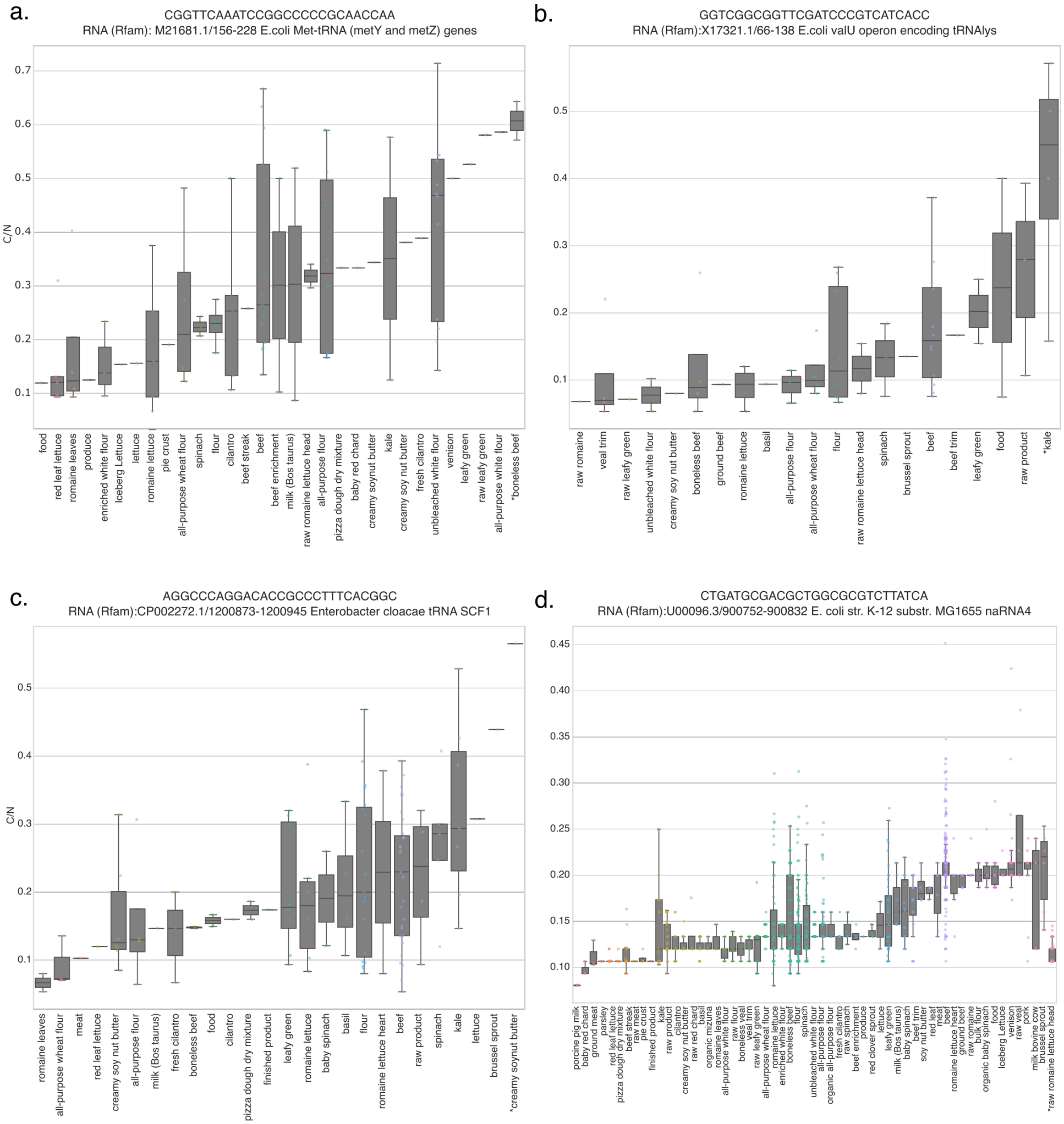


**Figure S11: Non-coding RNA show different target variability as a function of the isolation source in *E. coli* isolates. a.** Anchor CGGTTCAAATCCGGCCCCCGCAACCAA, mapping to a tRNA gene (Rfam: M21681.1/156-228), shows approximately a 6-fold difference in target diversity between isolates obtained from boneless beef compared to that of red leaf lettuce. **b.** Anchor GGTCGGCGGTTCGATCCCGTCATCACC, mapping to a tRNA gene (Rfam: X17321.1/66-138), shows approximately a 7-fold difference in target diversity between isolates obtained from kale to that of raw romaine. **c.** Anchor AGGCCCAGGACACCGCCCTTTCACGGC, mapping to a tRNA gene (Rfam: CP002272.1/1200873-1200945), shows approximately a 10-fold difference in target diversity between isolates obtained from romaine leaves to that of soynut butter. **d.** Anchor CTGATGCGACGCTGGCGCGTCTTATCA, mapping to a naRNA4 gene (Rfam: U00096.3/900752-900832), shows approximately a 5-fold difference in target diversity between isolates obtained from porcine pig milk to that of brussel sprouts.
